# Experience-independent transformation of single-cell 3D genome structure and transcriptome during postnatal development of the mammalian brain

**DOI:** 10.1101/2020.04.02.022657

**Authors:** Longzhi Tan, Wenping Ma, Honggui Wu, Yinghui Zheng, Dong Xing, Ritchie Chen, Xiang Li, Nicholas Daley, Karl Deisseroth, X. Sunney Xie

## Abstract

Both transcription and 3D organization of the mammalian genome play critical roles in neurodevelopment and its disorders. However, 3D genome structures of single brain cells have not been solved; little is known about the dynamics of single-cell transcriptome and 3D genome after birth. Here we generate a transcriptome atlas of 3,517 cells and a 3D genome atlas of 3,646 cells from the developing mouse cortex and hippocampus, using our high-resolution MALBAC-DT and Dip-C methods. In adults, 3D genome “structure types” delineate all major cell types, with high correlation between A/B compartments and gene expression. During development, both transcriptome and 3D genome are extensively transformed in the first postnatal month. In neurons, 3D genome is rewired across multiple scales, correlated with gene expression modules and independent of sensory experience. Finally, we examine allele-specific structure of imprinted genes, revealing local and chromosome-wide differences. These findings uncover a previously unknown dimension of neurodevelopment.

**HIGHLIGHTS:**

- Transcriptomes and 3D genome structures of single brain cells (both neurons and glia) in the developing mouse forebrain
- Cell type identity encoded in the 3D wiring of the mammalian genome (“structure types”)
- Major transformation of both transcriptome and 3D genome during the first month of life, independent of sensory experience
- Allele-specific 3D structure at 7 imprinted gene loci, including one that spans a whole chromosome

## INTRODUCTION

Two intimately related dimensions of the mammalian genome—gene transcription and three-dimensional (3D) genome architecture—are both crucial for the development of nervous systems. The dynamic interplay between cell-type-specific gene expression, chromatin structure, sensory experience, and other factors (e.g. epigenetic marks) underlies our brain’s immense plasticity and functions. Dysregulation of transcription and chromatin structure leads to debilitating neurodevelopmental disorders, including autism spectrum disorder, intellectual disability, and schizophrenia. For example, half of the top 102 autism-implicated genes are involved in gene transcription and chromatin organization (Satterstrom et al., 2020). Genome topology—in particular chromatin A/B compartments—also determines the distribution of somatic DNA damage and mutations in human neurons (Zhu et al., 2019), further shaping the genomic landscape of development, neurodegeneration, and cancers. It is therefore crucial to understand how the mammalian genome folds in 3D during development, and how it relates to the brain’s ever-changing transcriptome and diverse cell types.

Recent progress has been made to characterize transcriptome and 3D genome in the brain; however, existing methods and data have two major limitations. First, 3D genome structures of single brain cells have not been resolved. For example, bulk chromatin conformation capture (3C/Hi-C) assays can only measure certain statistical properties of 3D genome (i.e. population-averaged chromatin contact maps), but not actual 3D structures. These methods also lump together the diverse sub-types of neurons (and in many cases, glia, too) and mask critical cell-type-specific features (Lu et al., 2020; Won et al., 2016). So far, bulk Hi-C only revealed 3D genome refolding in neurons that are differentiated *in vitro* (Bonev et al., 2017; Lu et al., 2020; Rajarajan et al., 2018), or isolated at a single time point from the embryonic (day 14.5) mouse brain (Bonev et al., 2017). Single-cell 3C/Hi-C has been achieved in adult human brains (Lee et al., 2019), but had low resolution, yielded no 3D structures, and could not distinguish different neuronal sub-types—such as excitatory and inhibitory neurons, whose imbalance leads to many psychiatric disorders—based on structural information alone.

Second, the postnatal dynamics of single-cell transcriptome has not been comprehensively studied in the mammalian brain. Although transcriptional diversity has been extensively characterized in the adult brain (Tasic et al., 2016; Tasic et al., 2018; Zeisel et al., 2018) and more recently in the embryonic brain (Di Bella et al., 2020; La Manno et al., 2020), only a few studies have probed its postnatal dynamics, each with a limited set of time points (in mice): postnatal days (P) 2 and 11 for the whole brain (Rosenberg et al., 2018), P0, 4, 7, and 10 for the cerebellum (Carter et al., 2018), P1, 7, 90 for dopamine neurons (Tiklova et al., 2019), P10, 15, and 20 for the auditory cortex (Kalish et al., 2020), and P4, 8, 14, and 45 for the hypothalamus (Kim et al., 2020). Furthermore, most of these studies used low-sensitivity (split-pool or droplet-based) methods, which can only detect a small number of transcripts per cell and thus compromise data quality. A complete picture of postnatal gene expression in the mammalian brain is still lacking.

For the above reasons, existing studies can neither address the true relationship between the brain’s diverse transcriptional cell types and their underlying 3D genome “structure types”, nor trace their dynamic interplay during development *in vivo*.

Here we fill in this longstanding knowledge gap by providing a high-resolution database of both single-cell transcriptomes and single-cell 3D genome structures throughout postnatal brain development. Using our latest, highly sensitive and accurate multiple annealing and looping based amplification cycles for digital transcriptomics (MALBAC-DT) method (Chapman et al., 2020), we first generated a transcriptome atlas of the developing mouse forebrain, encompassing ∼3,500 single cells (as isolated nuclei) from 2 brain regions and 7 time points across the first 6 months of life. Using a fully streamlined version of our diploid chromatin conformation capture (Dip-C) method (Tan et al., 2018; Tan et al., 2019), we further generated a comprehensive 3D genome atlas, encompassing ∼2,000 single-cell contact maps and ∼800 high-resolution 3D structures from 2 brain regions and 6 time points across the first year of life. Finally, to determine whether 3D genome change is genetically predetermined or depends on early-life experience, we analyzed 3D genome in the visual cortex of sensory-deprived and control mice, with ∼1,700 single cells sampled weekly (5 time points) during the first postnatal month.

With our uniquely high sensitivity and accuracy (for transcriptome) and high spatial resolution (for 3D genome: 20 kb, or ∼100 nm), we directly addressed several fundamental questions in developmental biology. On the transcriptome side, we observed a major transformation around P14 in all 3 main cell lineages: neurons, astrocytes, and oligodendrocytes. In neurons, despite clear brain-region and cell-type specialization (producing 16 sub-types in adults), the primary source of transcriptome variation was explained by two developmentally regulated, correlated gene modules—which we termed “neonatal” and “adult” modules.

On the 3D genome side, we found that cell type identity was encoded in the 3D wiring of the genome, because 3D genome structure alone could separate single cells into 13 neuron and glia types, which we termed “structure types”. Between structural types, differential chromatin A/B compartmentalization correlated well with cell-type-specific gene expression. We observed a major 3D genome transformation between P7 and 28 in all 3 cell lineages, coincident with their transcriptome transformation. Most notably, the 6 adult neuron sub-types only emerged during this transition. Neonatal neurons, in contrast, adopted a more primordial 3D genome state that resembled embryonic neurons and neurons differentiated *in vitro* (Bonev et al., 2017).

We gained mechanistic insights into this transformation by analyzing 3D structures across multiple spatial scales. With Dip-C’s unique capability, we revealed large-scale, neuron-specific radial reconfiguration: inward movement of many regions across the genome—including the majority of mouse Chr 7, 17, and 19—that preferentially harbored clustered gene families such as olfactory receptors (ORs). This phenomenon could be linked to global non-CpG DNA methylation during the same developmental window (Lister et al., 2013), and shared some striking similarity (albeit with important differences) with 3D remodeling of a peripheral nervous system (Clowney et al., 2012; Monahan et al., 2019; Tan et al., 2019). Although not directly correlated with gene expression modules, such radial movement might help to further silence OR expression (Winick-Ng et al., 2020).

We also observed local and long-range 3D rewiring of many developmental and/or disease-implicated genes and their enhancers. Changes in single-cell chromatin A/B compartments (scA/B) correlated with developmentally regulated gene modules, although many genes displayed discordant (or temporally shifted) changes in transcriptome and 3D genome.

To functionally interrogate 3D genome transformation, we reared mice in complete darkness from birth. This sensory deprivation had little effect on 3D genome in the visual cortex, suggesting that our observed transformation was genetically predetermined, rather than experience-induced.

Finally, we report a genome-wide, allele-specific survey of imprinted genes. We found parent-of-origin-specific structure for at least 7 of the 29 known imprinted loci (Perez et al., 2015). In contrast, other genomic regions rarely adopted allele-specific structure. In one extreme case, allelic difference could be seen extending tens of Mbs—from the Prader-Willi/Angelman syndrome (PWS/AS) locus to nearly the entire mouse Chr 7.

In summary, this work offers both a rich resource for exciting discoveries in mammalian neurodevelopment, and a new experimental and analytical paradigm for multi-omic, single-cell 3D genome analyses that is widely applicable to many developmental processes, such as organogenesis, cancers, and aging.

## RESULTS

### Our highly sensitive and accurate MALBAC-DT method enables a single-cell transcriptome atlas of postnatal brain development

To create a transcriptome atlas of the developing mouse forebrain, we performed our high-precision single-cell RNA-seq method, termed MALBAC-DT (Chapman et al., 2020), on 3,517 single cells from the cortex and hippocampus at 7 different ages: on P1, P7, P14, P21, P28, P56, and P180 (Figure 1A top, Figure 1B top). Dense sampling ensured weekly resolution in the first postnatal month—a critical period for cognitive plasticity, in which the brain undergoes numerous molecular and functional changes. To ensure uniformity, all animals were males from the widely used inbred strain B6 (here the C57BL/6N sub-strain). To minimize stress (Lacar et al., 2016), single cells were rapidly isolated from dissected tissues as individual nuclei (Krishnaswami et al., 2016) (Methods). Note that we still referred to them as “cells.”

**Figure 1.**
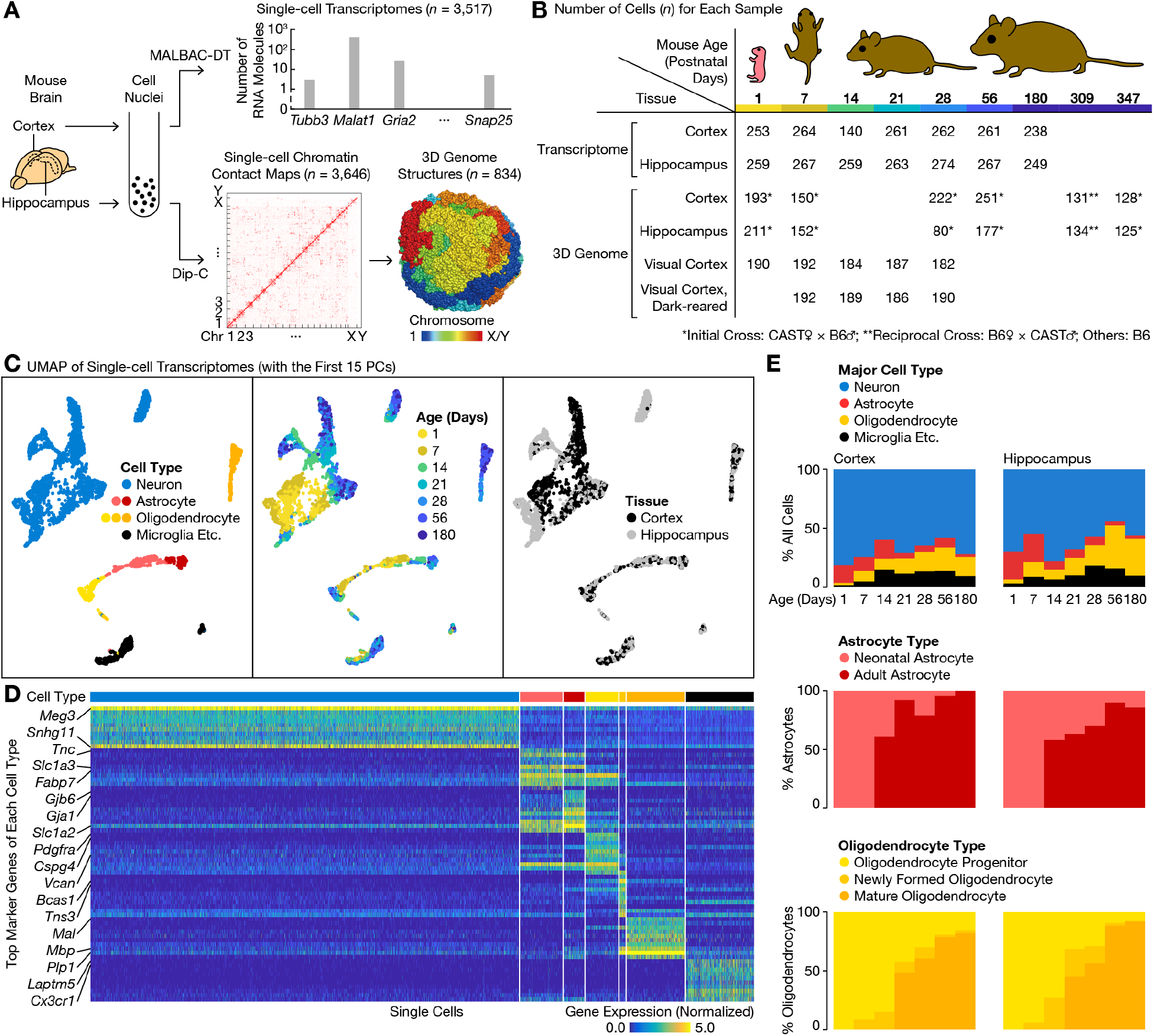
A single-cell transcriptome atlas of the mouse forebrain during postnatal development with MALBAC-DT. (A) Schematic of the study. (B) Number of single cells for each sample. (C) UMAP of single-cell transcriptomes. UMAP was performed on the first 15 PCs of normalized gene expression levels (with SCTransform, which used the top 3,000 variable genes by default). (D) Expression of top cell-type marker genes among single cells. For each cell type, the top 10 marker genes were identified with Seurat. Expression levels were normalized with SCTransform in Seurat. (E) Composition of transcriptome types for each tissue and age. For all cells (top), *n* was shown in (B). For astrocytes (middle), *n* = 38, 31, 23, 13, 14, 22, 6, 62, 62, 19, 19, 20, 10, 7 from left to right. For oligodendrocytes (bottom), *n* = 5, 23, 13, 33, 43, 51, 38, 8, 34, 22, 38, 48, 97, 77.

MALBAC-DT allowed us to measure single-cell transcriptome with unprecedented sensitivity and accuracy (Chapman et al., 2020). We detected an average of 12.8 k mRNA molecules (also known as unique molecule identifiers (UMIs); SD = 8.4 k, minimum = 1.7 k, maximum = 49.8 k) and 3.4 k genes (SD = 1.3 k, minimum = 1.0 k, maximum = 7.8 k) per cell (Figure S1). This was an order of magnitude higher than any previous studies that probed part of postnatal cortical development in mice: 1.0 k UMIs and 0.7 k genes with SPLiT-seq (Rosenberg et al., 2018), and 1.9 k UMIs and 1.2 k genes with inDrops (Kalish et al., 2020). MALBAC-DT was also more accurate, because it avoided a UMI-overcounting issue that plagued most other methods (Chapman et al., 2020). Our data therefore provided the most comprehensive temporal coverage (P1 to 180) and the highest per-cell data quality to date.

### Transcriptome behind all 3 main lineages (neurons, astrocytes, and oligodendrocytes) substantially changes around P14

To visualize transcriptional diversity and dynamics among cells, we first performed uniform manifold approximation and projection (UMAP) on the first 15 principal components (PCs) of single-cell transcriptomes, using Seurat v3 (Stuart et al., 2019) and its latest normalization method SCTransform (Hafemeister and Satija, 2019). Cells clearly separated into distinct clusters, each with unique age and tissue patterns (Figure 1C). We referred to these cell-type clusters as “transcriptome types” to distinguish from 3D genome analysis in later sections.

Louvain clustering of single-cell transcriptomes—based on the first 15 PCs—identified 4 major cell types: neurons, astrocytes, oligodendrocytes, and other cells (“microglia etc.”: containing primarily microglia and a small number of vascular cells) (Figure 1C left, Figure 1D, Figure 1E top, Figure S2). Expression patterns of known marker genes were shown in Figure S2. All our identified markers were shown in Table S1.

A fundamental question in developmental biology is to what extent postnatal cells continue to mature in their gene expression profile. This is especially important for neurons in the brain— most of which have become postmitotic at this point, and must last a lifetime. On our UMAP plot, interestingly, within each major type, cells tended to segregate primarily by age, with the clearest visual distinction between P7 and 14 (Figure 1C middle). This suggested a major transcriptome transformation after birth, in all cell lineages of the brain.

We first examined oligodendrocytes and astrocytes. Oligodendrocytes segregated into 3 distinct sub-types: oligodendrocyte progenitors (*Pdgfra*+), newly formed oligodendrocytes (*Tns3*+), and mature oligodendrocyte (*Mal*+) (Figure 1C left), consistent with a previous single-cell study of the adult visual cortex (Tasic et al., 2016) and a bulk microarray study of the forebrain (Cahoy et al., 2008). In both brain regions, starting from a single transcriptome type (oligodendrocyte progenitors) on P1, postnatal development led to the emergence of two new types over time: newly formed oligodendrocytes on P7 and mature oligodendrocytes on P21, the latter of which later dominated adult populations (Figure 1E bottom).

Similarly, transcriptome transformation in astrocytes led to the emergence of a more mature type (“adult astrocytes”; *Gjb6*+) on P14 in addition to the neonatal type (*Tnc*+) (Figure 1E middle), consistent with a bulk microarray study comparing P1–8 and P17–30 astrocytes in the forebrain (Cahoy et al., 2008).

### Neurons consist of complex transcriptome types with distinct temporal, regional, and functional specificities

We now focus on neurons—the most prevalent lineage in both brain regions. Different functional sub-types of neurons, such as excitatory and inhibitory neurons, are known to carry out different computations. Neurons in different brain regions, and/or at different developmental stages, can also exhibit different properties.

To investigate the transcriptomic basis of such enormous diversity, we isolated all 2,277 neurons and performed another round of normalization, followed by UMAP on the first 30 newly generated PCs (Figure 2A). This procedure ensured that genes most relevant for neuronal diversity were selected during normalization, without interference from other cells.

**Figure 2.**
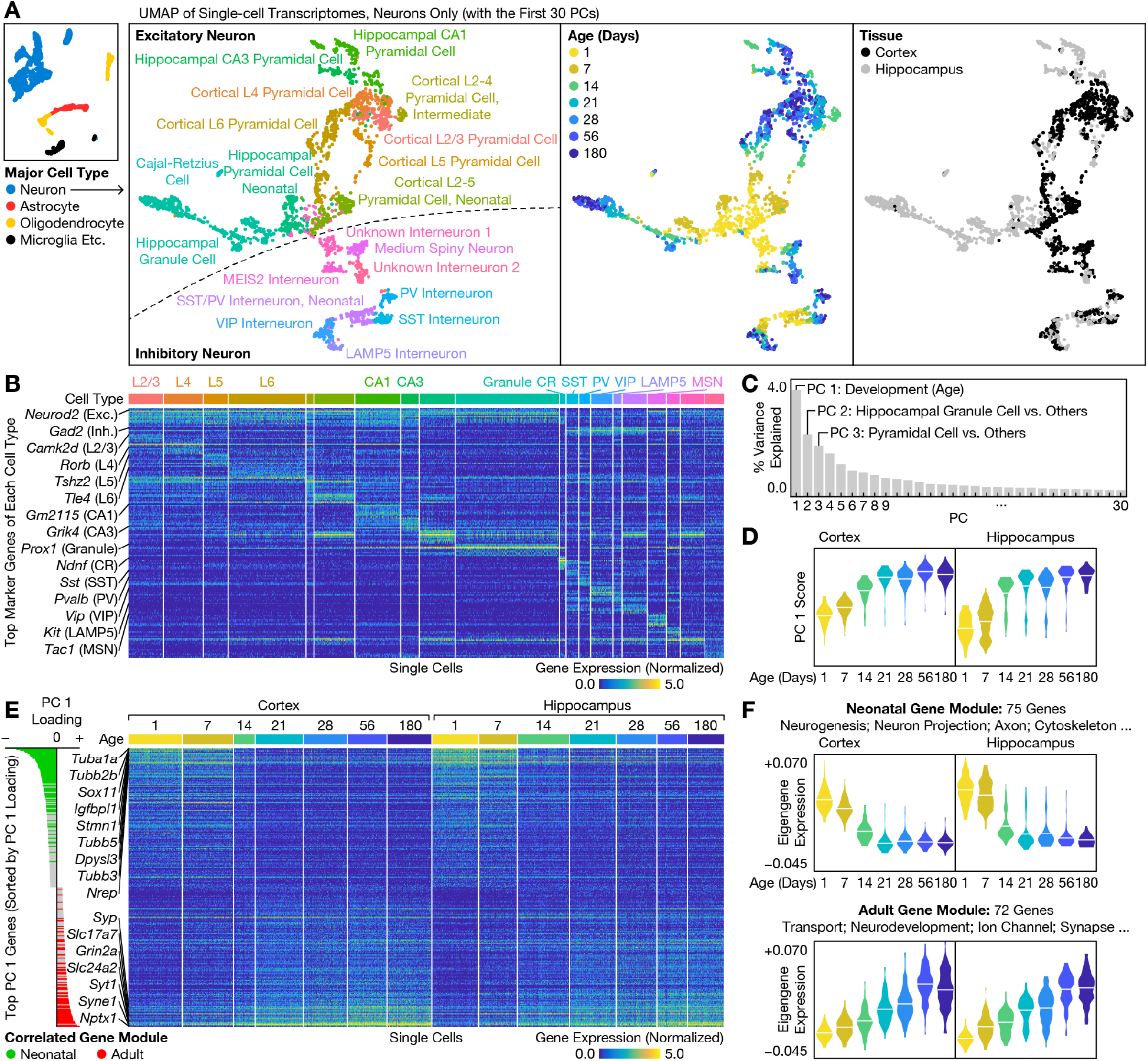
Transcriptome variation among neurons. (A) UMAP of single-cell transcriptome among neurons. UMAP was performed on the first 30 PCs (by default) of normalized gene expression levels (neurons only). (B) Expression of top cell-type marker genes among single neurons. For each cell type, the top 20 marker genes were identified with Seurat. If a gene was identified for multiple cell types, only the one with the highest fold change was kept. (C) Percent variance explained by each PC. (D) Distribution of single neurons along PC 1 for each tissue and age. Horizontal lines denoted the mean. *n* = 205, 196, 83, 184, 169, 151, 171, 180, 147, 201, 178, 156, 117, 139 from left to right. Bandwidth was 1.5. Axes limits were −40 and +25. (E) Expression of top PC 1 genes among single neurons. Axes limits for PC 1 loading (left) were −0.220 and +0.098. Top 150 genes on each side were shown. Correlated gene modules were identified with WGCNA. (F) Eigengene expression of the 2 correlated gene modules, among single neurons for each age and tissue. Selected key words of enriched GO terms were shown. Horizontal lines denoted the mean. *n* was the same as in (D). Bandwidth was 0.003.

On the UMAP plot, neurons formed clusters with a complex pattern according to their ages, brain regions, and functions (Figure 2A). For example, the two brain regions (cortex and hippocampus) were largely separated among excitatory neurons, yet relatively mixed among inhibitory neurons (Figure 2A right). For most sub-types, a prominent age gradient could be seen—usually with the clearest visual separation between P7 and 14 (Figure 2A second to the right), consistent with the UMAP plot of all cells (Figure 1C middle).

Louvain clustering classified neurons into 20 transcriptome types (Figure 2A second to the left, Figure 2B). Because temporal dynamics was analyzed in the next section, here we merged different developmental stages of the same sub-type when possible, only keeping immature types (named “neonatal” or “intermediate”) when necessary. This procedure led to 16 main types and 4 immature types.

Among the 16 main types, 8 were excitatory: 4 cortical pyramidal cell types based on their cortical layers (L2/3: *Camk2d*+; L4: *Rorb*+; L5: *Tshz2*+; L6: *Tle4*+, among which a small sub-cluster of *Ctgf*+ L6b cells were visible), 2 hippocampal pyramidal cell types of the *Cornu Ammonis* (CA) region (CA1: *Gm2115*+; CA3: *Grik4*+), hippocampal granule cells of the dentate gyrus (DG) region (*Prox1*+), and Cajal–Retzius (CR) cells (*Nhlh2*+, also with much higher expression of *Ndnf* and *Reln* than in inhibitory neurons) (Figure 1A, Figure 1B). Expression patterns of known marker genes were shown in Figure S3. All our identified markers were shown in Table S2.

The other 8 were inhibitory: 5 known interneuron types (SST: *Sst*+; PV: *Pvalb*+; VIP: *Vip*+; LAMP5: *Lamp5*+, named as in (Tasic et al., 2018), some cells of which are called NDNF interneurons by (Tasic et al., 2016) and others; MEIS2: higher expression of *Meis2* than in other neurons, named as in (Di Bella et al., 2020; Jin et al., 2019; Tasic et al., 2018)), medium spiny neurons (MSN: *Tac1*+, also *Meis2+* but lower than in MEIS2 interneurons; also seen in (Jin et al., 2019)), and 2 unknown interneuron types (Figure 1A, Figure 1B, Figure S3).

The remaining 4 types were immature neurons that corresponded to multiple adult types: neonatal cortical L2–5 pyramidal cells (mostly from P1 and 7), intermediate cortical L2–4 pyramidal cells (P14), neonatal hippocampal pyramidal cells (P1 and 7), and neonatal SST/PV interneurons (P1 and 7) (Figure 1A, Figure 1B, Figure S3).

### Two developmentally regulated gene modules (“neonatal” and “adult”) explain the top component of transcriptome variation among neurons

Given the extreme complexity of transcriptome among neurons, a critical question is what the primary sources of variability are. We used principal component analysis (PCA) to dissect the distinct (mutually orthogonal) sources of cell-to-cell heterogeneity (Figure 2C). Despite neurons’ well-known transcriptional specialization based on functions and regions, the first PC— explaining 3.6% of the total variance (of the top 3,000 variable genes in neurons, as selected by SCTransform)—corresponded to developmental progression at different ages. In comparison, the next two PCs, which distinguished major sub-types (granule and pyramidal cells, respectively), explained 2.1% and 1.7% of the total variance.

Neurons were positioned along the first PC primarily based on age, with cells from P1 and 7 spread out on the negative side, P21 and above tightly concentrated on the positive end, and P14 in between (Figure 2D). Note that the hippocampus harbored more adult cells on the negative side, partly as a result of adult neurogenesis of DG granule cells (Figure 2A).

To identify genes underlying transcriptome maturation along this PC, we examined genes with the highest PC 1 loadings (in absolution values; Figure 2E, Table S3, Table S4). As expected, genes with the most negative loadings were primarily expressed by neonatal neurons, while genes with the most positive loadings were more expressed by adult neurons (Figure 2E).

We identified correlated gene modules with WGCNA (Langfelder and Horvath, 2008) as an independent confirmation. The 2 largest modules—which we termed the “neonatal” and “adult” modules—corresponded well with PC 1 loadings. Among the 3,000 genes that were used in PCA, the 75 genes in the neonatal module ranked between the 1st and the 123rd most negative (mean rank = 44), while the 72 genes in the adult module ranked between the 1st and the 148th most positive (mean rank = 53) (Figure 2D left, Table S5). Note that for maximal confidence, the above modules were identified from the top 3,000 variable genes in neurons, as selected by SCTransform. A less stringent list from the top 10,000 variable genes—yielding 122 genes in the neonatal module and 151 in the adult one—was shown in Table S6.

We visualized the expression patterns of the 2 correlated gene modules with their eigengenes—a weighted average of individual genes in a module, as produced by WGCNA (Langfelder and Horvath, 2008). The neonatal eigengene was highly expressed on P1 and 7, sharply declined on P14, and remained low from P21 onward (Figure 2F top). In contrast, the adult eigengene steadily increased over time until plateauing around P56 (Figure 2F bottom). Expression patterns of all individual genes were shown in Figure S4. These two modules therefore represented two distinct modes of transcriptome transformation in postnatal neurons.

### Genes in the two developmental modules carry out crucial neuronal functions

Gene ontology (GO) analysis with PANTHER (Thomas et al., 2003) revealed enriched gene functions among the 2 correlated modules. Genes in the neonatal modules were enriched in neurogenesis (36 genes; FDR = 6 × 10^−17^), cellular developmental process (43; 4 × 10^−13^), neuron projection development (22; 7 × 10^−13^), axon development (17; 8 × 10^−12^), structural constituent of cytoskeleton (5; 4 × 10^−3^), and polymeric cytoskeletal fiber (12; 4 × 10^−4^). A full list of enriched GO terms was shown in Table S5 and Table S6. Neonatal genes included cytoskeleton genes (e.g. *Tuba1a/b2a/b2b/b3/b5, Stmn1/2, Syne2*), axon guidance molecules (e.g. *Gap43, Nrp1, Dpysl3/5, Plxnb1, Sema3c*), and transcription factors (e.g. *Lhx2, Sox4/11, Tcf4, Nfib, Nrep, Rcor2, Ctnnb1*). Some of the functional enrichments were consistent with a bulk microarray study of the visual cortex (whole tissue) at P0, 14, 28, and 60 (Lyckman et al., 2008).

Genes in the adult module were enriched in transport (32 genes; FDR = 7 × 10^−6^), nervous system development (24; 5 × 10^−5^), regulation of signaling (29; 5 × 10^−5^), regulation of synaptic plasticity (9; 5 × 10^−5^), ion channel activity (13; 9 × 10^−7^), and synapse (30; 8 × 10^−15^). Adult genes included ion channels (e.g. *Grin2a, Ano3, Gabra1/d, Kcna2/j9/k9/ip3, Slc17a7/24a2*), synaptic proteins (e.g. *Syt1/13, Syp, Nptx1, Lamp5*), and cytoskeleton genes (e.g. *Syne1, Nefl/m, Map1a*). Similarly, some of the enrichments were consistent with a whole tissue study (Lyckman et al., 2008).

Most genes in the 2 modules were shared by all neuron sub-types examined—some genes further shared by neurons in the peripheral nervous system (for example, olfactory sensory neurons (OSNs) that we previously studied (Magklara et al., 2011; Tan et al., 2015; Tan et al., 2019)); however, some other genes were sub-type-specific (Figure S4). For example, in the neonatal module, *Sema3c, Igfbpl1*, and *Zbtb20* were specific to the hippocampus. In the adult module, *Slc17a7* was specific to excitatory neurons (except CR cells).

Note that a few genes were not primarily expressed by neurons. For example, *Vcan* in the neonatal module was highly expressed by oligodendrocyte progenitors, while *Mbp* in the adult module was highly expressed by mature oligodendrocyte. It was unclear whether their neuronal expression arose from diffusing RNA molecules during cell isolation. Their non-zero neuronal expression was also detected by other datasets (Zeisel et al., 2018).

### A streamlined Dip-C method resolves 3D genome structures of single brain cells across postnatal development

To understand the structural basis of the observed transcriptional transformation, we now turn to 3D genome architecture. Using our high-resolution single-cell 3C/Hi-C method, termed Dip-C (Tan et al., 2018), we have previously solved the 3D genome structures of single sensory neurons in the mouse eye and nose, characterizing their reorganization during postnatal development (Tan et al., 2019).

Here we further optimized Dip-C for rapid, high-throughput sample processing, by enabling the use of multi-channel pipettes at all steps, and handling and indexing 4–8 96-well plates at once (Methods). Furthermore, the streamlined protocol can now be performed with commercially available, low-cost reagents only, and is potentially compatible with full automation.

To create a 3D genome atlas of the developing mouse forebrain, we performed the streamlined Dip-C workflow on 1,954 single cells from the cortex and hippocampus at 6 different ages throughout the first postnatal year: on P1, P7, P28, P56, P309, and P347 (Figure 1A bottom, Figure 1B middle). To enable 3D genome reconstruction (by resolving two parental haplotypes), all animals were filial 1 (F1) hybrids from a CAST/EiJ♀ × C57BL/6J♂ cross (denoted the initial cross) except the P309 animal, which was from a reciprocal cross (C57BL/6J♀ × CAST/EiJ♂) (Figure 1B) for parent-of-origin analysis (see the last section). Single cells were rapidly isolated as individual nuclei (Krishnaswami et al., 2016; Lacar et al., 2016) and chemically fixed in a modified buffer that preserved nuclear morphology (Methods).

We obtained an average of 401 k chromatin contacts per cell (SD = 116 k, minimum = 101 k, maximum = 1.20 m) (Figure S5, Table S7). Among the cells, 834 (43%) yielded high-quality 3D structures at 20-kb resolution, defined as root-mean-square deviation (RMSD) ≤ 1.5 particle radii (average = 0.90, SD = 0.28 among those cells; each 20-kb particle has a radius of ∼100 nm) (Table S7). These high-resolution contact maps and 3D structures provided a unique opportunity to study cell-type-specific genome structure, its developmental time course, and its relationship with gene expression.

### 3D genome structure alone delineates 13 major neuronal and glial types

A fundamental question in cell biology is whether each cell type/sub-type corresponds to an underlying 3D genome structure type, and if so, when these structure types emerge during development. We have previously used PCA of single-cell A/B compartment (scA/B) values— approximately defined for each genomic locus as the rank-normalized CpG density in its 3D neighborhood—to distinguish cell types in the human blood (Tan et al., 2018) and in the mouse eye and nose (Tan et al., 2019), a procedure which we termed “structure typing”.

However, cell types are more complex in the brain, where related cell sub-types can be challenging to resolve. For example, a recent, low-resolution single-cell 3C/Hi-C study could not tell apart single neurons of different sub-types from contact maps alone (Lee et al., 2019). To solve this problem, we developed a new pipeline, which performed t-distributed stochastic neighbor embedding (t-SNE) on the first 20 PCs of the scA/B matrix (1-Mb bins, therefore ∼3,000 values per cell) (Figure 3A) in a manner similar to those widely used in transcriptome studies. This approach clearly visualized multiple clusters of cells, which we termed “structure type”, each with distinct age and tissue profiles (Figure 3A).

**Figure 3.**
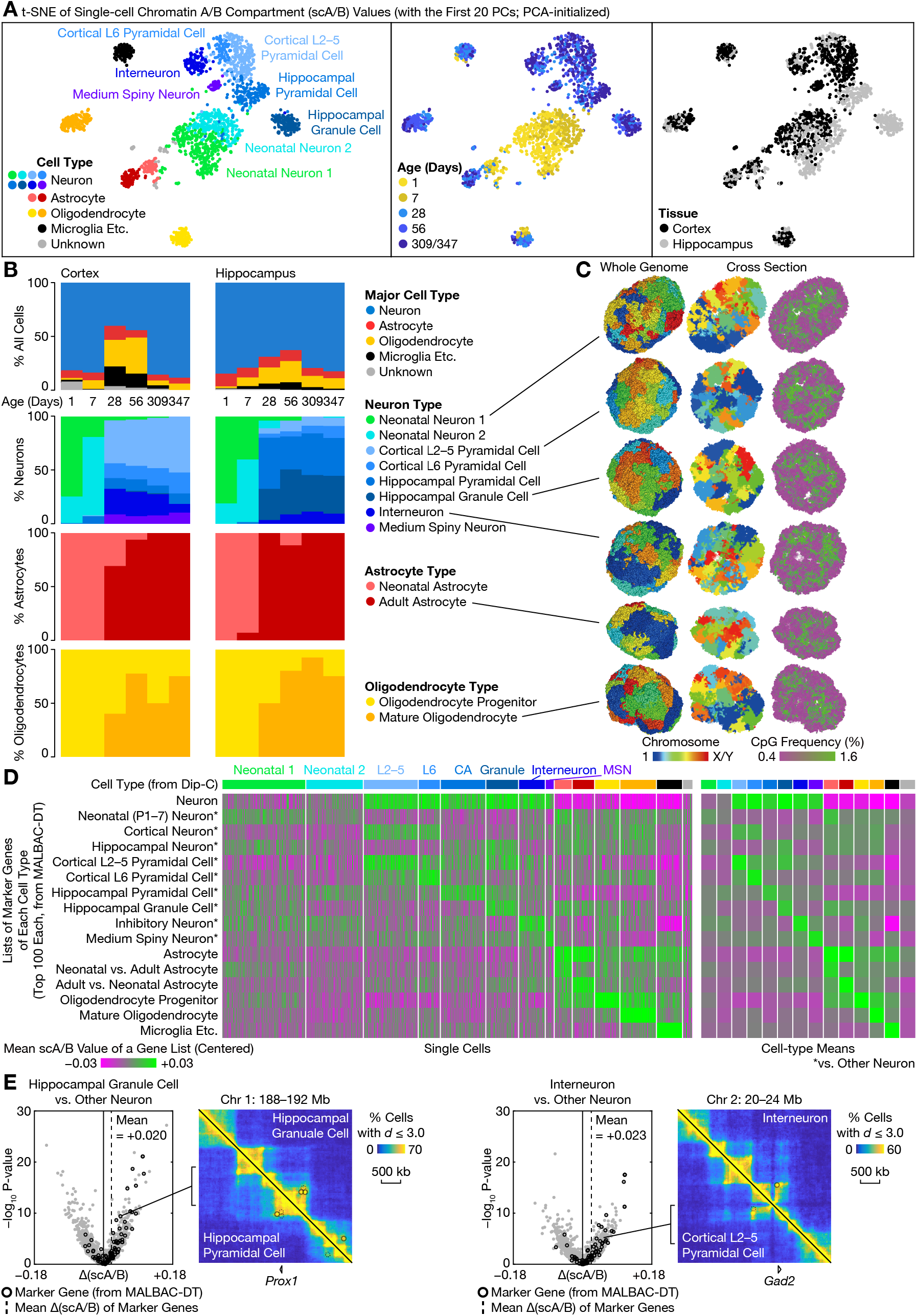
A single-cell 3D genome atlas of the mouse forebrain during postnatal development with Dip-C. (A) t-SNE of single-cell 3D genomes. t-SNE was performed on the first 20 PCs of scA/B values. (B) Composition of structure types for each tissue and age. Note that the decrease in glia at the 2 oldest ages might be caused by batch difference in filtration during nuclei isolation. For all cells (top), *n* was shown in Figure 1B. For neurons (second row), *n* = 157, 125, 89, 111, 112, 113, 178, 120, 55, 111, 106, 103 from left to right. For astrocytes (third row): *n* = 30, 12, 38, 23, 9, 7, 28, 16, 9, 19, 10, 8. For oligodendrocytes (bottom): *n* = 3, 12, 55, 84, 6, 8, 4, 12, 12, 35, 14, 12. (C) Example 3D genome structures. (D) Mean scA/B values of lists of cell-type marker genes among single cells (left), and averaged for each structure type (right). For each transcriptome type, the top 100 marker genes were identified from MALBAC-DT data with Seurat. For the number of cells in each structure type, *n* = 346, 239, 228, 94, 189, 136, 111, 37, 75, 93, 105, 152, 108, 41 from left to right. (E) Example volcano plots of differential scA/B between cell types, with 3D proximity maps (percent cells where two given 20-kb particles were within a 3D distance of 3.0 particle radii) for example genes with both cell-type-specific expression and scA/B. For volcano plots, P-values were from two-sided Mann–Whitney U tests; *n* = 136 vs. 1,244 (left) and 111 vs. 1,269 (right). For 3D proximity maps, circles denoted differential chromatin loops; *n* = 86 vs. 624 cells × 2 alleles (left) and 57 vs. 653 cells × 2 alleles (right). Note that some regions (for example, near *Gad2*) were unmappable, leading to poor estimation of 3D distances.

Structure typing was robust against our choice of parameters. Visualization of structure types was relatively unchanged when contacts were down-sampled to 100 k per cell (or only moderately degrades at 50 k per cell), when the genome was binned every 100 kb, when fewer (10) or more (50) PCs were retained, or when t-SNE was substituted by UMAP (Figure S6).

Hierarchical clustering of cells—based on the first 20 PCs of the scA/B matrix—identified 13 structure types and 3 smaller, unknown clusters (Figure 3A left). By multi-modal integration with our MALBAC-DT data and published omic data (see below), we were able to identify all 13 structure types: 8 neuron types, 2 astrocyte types, 2 oligodendrocyte types, and 1 other type (“microglia etc.”) (Figure 3A).

### “Structure types” are highly correlated with their corresponding transcriptome types

A major unresolved question about single-cell 3D genome structure is to what extent it corresponds to single-cell transcriptome and other “omes.” We have previously shown in a few examples that cell-type-specific genes, when averaged, tend to exhibit higher scA/B values (namely, more A-like and more euchromatic) specifically in single cells of the corresponding cell type (Tan et al., 2019). However, a systematic investigation—especially in a complex organ like the brain—has not be conducted.

Our matched MALBAC-DT and Dip-C datasets provided a unique opportunity to interrogate the relationship between structure and transcriptome types. Here we developed a new pipeline to compare cell-type-specific gene expression from single-cell transcriptome data and differentially structured regions from single-cell 3D genome data (Methods).

We matched each structure type to its corresponding transcriptome type(s), by first calculating the mean scA/B value for each set of cell-type marker genes (identified from MALBAC-DT data) in each Dip-C cell (Figure 3D). For each gene set, scA/B values typically showed specific enrichment in one or a few structure types. Patterns of scA/B values on the t-SNE plot were shown in Figure S7. This high correlation enabled unambiguous cell-type identification of all 13 structure types.

For example, the top 100 microglia-specific marker genes from MALBAC-DT had a mean scA/B value (on a 0–1 scale) of 0.727 (SD = 0.012, SEM = 0.001) among Dip-C cells in the microglia structure type—0.060 higher than the next highest structure type (mature oligodendrocytes: mean = 0.667, SD = 0.013, SEM = 0.001) (Figure 3D last row). This procedure thus positively identified the microglia structure type.

The 2 oligodendrocyte structure types were similarly identified as oligodendrocyte progenitors (with a 0.029 scA/B margin between the best match and the next) and mature oligodendrocytes (0.043), respectively. However, the intermediate transcriptome type—newly formed oligodendrocytes—matched equally well to both structure types (only a 0.001 margin between the best match and the next, compared to a 0.015 margin to the 3rd best). This might represent a mismatch between structural and transcriptional dynamics during a transitional stage of oligodendrocyte maturation. Alternatively, our Dip-C dataset might lack resolution or sample size.

Although marker gene sets for all, neonatal, or adult astrocytes each matched equally well to the 2 astrocyte structure types, a more nuanced analysis using discriminative marker genes (between neonatal and adult astrocytes) successfully distinguished the 2 (0.013 margin for neonatal, and 0.021 for adult) (Figure 3D). The need for additional analysis might indicate high molecular similarity between the 2 astrocyte types.

The 8 neuron structure types were first identified as a whole, using neuron-specific marker genes (0.019 margin, or 0.044 if neonatal neurons were excluded) (Figure 3D first row). Each adult type was then individually identified using discriminative markers between different neuron sub-types (margins: cortical L2–5 pyramidal cells = 0.011, cortical L6 pyramidal cells = 0.018, hippocampal pyramidal cells = 0.009, hippocampal granule cells = 0.019, interneurons = 0.019, and MSNs = 0.017). Figure 3E showed cell-type-specific 3D genome around two example marker genes; in both cases, cell-type-specific chromatin loops (potentially enhancers) were also observed. Furthermore, among the 4 excitatory types, brain-region-specific marker genes corresponded well with brain-region difference in scA/B (Figure 3D). Finally, we named the two remaining types neonatal neurons 1 and 2, which primarily consisted of neurons from P1 and 7, respectively.

Similar to oligodendrocytes, we observed fewer neuron structure types (6 in adults) (Figure 2A) than transcriptome types (16 in adults) (Figure 3A). For example, we did not observe specific scA/B patterns for cortical L2/3 or L5 marker genes (although some pattern could be seen for L4 markers). Their finer structural difference might require a larger sample size to resolve.

As an additional confirmation of the high correlation between cell-type-specific gene expression and scA/B, we directly compared differentially A/B-compartmentalized regions in Dip-C to MALBAC-DT marker genes. In particular, for each structure type, we systematically identified top regions (and the genes that they harbored) with differential scA/B (Table S8). For example, our scA/B analysis covered 2,410 1-Mb bins across the mouse autosome; the top 100 granule-cell-specific marker genes corresponded to 98 such bins—66 (67%) of which had on average higher scA/B (regardless of statistical significance) in granule cells than in other neurons (Figure 3E). This represented a significant enrichment with respect to the whole genome (67 vs. 52%; two-tailed Fisher’s exact test: P = 9.7 × 10^−5^).

### Structure types correlate with other omic data

We further confirmed the cell-type identity of structure types by integrating published 3D genome, transcriptome, methylome, and chromatin accessibility data. We first projected all available bulk Hi-C data of the mouse brain onto our t-SNE plot (Figure S8). Bulk Hi-C of astrocytes differentiated *in vitro* (Zhang et al., 2014) projected to the center of our 2 astrocyte structure types, validating their cell-type assignments. Among neurons, bulk Hi-C of adult cortical neurons (Jiang et al., 2017) and of adult hippocampal neurons (Fernandez-Albert et al., 2019) projected to our cortical pyramidal cells and hippocampal granule cells, respectively. Interestingly, bulk Hi-C of embryonic (day 14.5) cortical neurons and of cortical neurons differentiated *in vitro* (Bonev et al., 2017) both projected next to our youngest neonatal neurons (primarily from P1). This indicated that neurons did not attain structural maturity when cultured and differentiated *in vitro*, which had important implications for stem cell biology.

We next looked at published transcriptome (bulk and single-cell) data (Figure S9, Table S9). Similar to our MALBAC-DT data, cell-type-specific marker genes from published data (Cahoy et al., 2008; Habib et al., 2016; Rosenberg et al., 2018; Zhang et al., 2014) matched the expected structure types. Our results were therefore robust against different single-cell isolation and RNA-seq methods.

Finally, we extended the correlation between 3D genome architecture and gene expression to other modalities—accessible chromatin regions and un-methylated DNA regions (Figure S9). For example, the top 11 k microglia-specific accessible regions (Cusanovich et al., 2018) showed the highest mean scA/B values in the microglia structure type (0.026 margin), while the top 17 k oligodendrocyte-specific accessible regions (Cusanovich et al., 2018) had the highest scA/B in the mature oligodendrocytes (0.012 margin). Among neurons, both the top 18 k SST-interneuron-specific accessible regions (Cusanovich et al., 2018) and the top 35 k SST-interneuron-specific un-methylated regions (Luo et al., 2017) matched our interneurons (both 0.003 margin), while the top 5 k granule-cell-specific accessible regions (Lareau et al., 2019) matched our granule cells (0.006 margin).

### Structure types underlying all 3 lineages (neurons, astrocytes, and oligodendrocytes) fundamentally change between P7 and 28

We now examine the temporal dynamics of the 13 structure types. Most notably, between P7 and 28, structure type composition shifted drastically and concomitantly for neurons, astrocytes, and oligodendrocytes, indicating a major transformation of the underlying 3D genome organization in all 3 lineages in the first postnatal month (Figure 3B)—coincident with their transcriptome transformation.

In particular, each cell lineage shifted from its neonatal structure types into adult ones between P7 and 28 (Figure 3B), mirroring their transcriptional changes (Figure 1E). In glia, this led to a change from neonatal to adult astrocytes, and from oligodendrocyte progenitors to mature oligodendrocytes.

Interestingly, among neurons, only adult cells (P28 onward) clearly separated into distinct sub-types on the t-SNE plot (Figure 3A). Neonatal neurons (P7 and earlier), on the other hand, are more intermixed on the plot (Figure 3A)—albeit already exhibiting early signs of structural specialization, such as moderate separation between the 2 brain regions (Figure 3A) and apparent clustering of some neonatal granule cells at the bottom corner (Figure S7). Note that because the majority of brain neurons neither divide nor die after birth, their structure and transcriptome types must be directly converted within each cell between P7 and 28.

To gain mechanistic insights into this developmental conversion of structure types, we ask how neonatal and adult structure types differ. On the largest scale, we looked at the distribution of contact distances among intra-chromosomal contacts, and the percentage of inter-chromosomal contacts (Figure 4A). In both astrocytes and oligodendrocytes, postnatal development led to an increase in very long-range contacts (10–100 Mb) (Figure 4A top).

**Figure 4.**
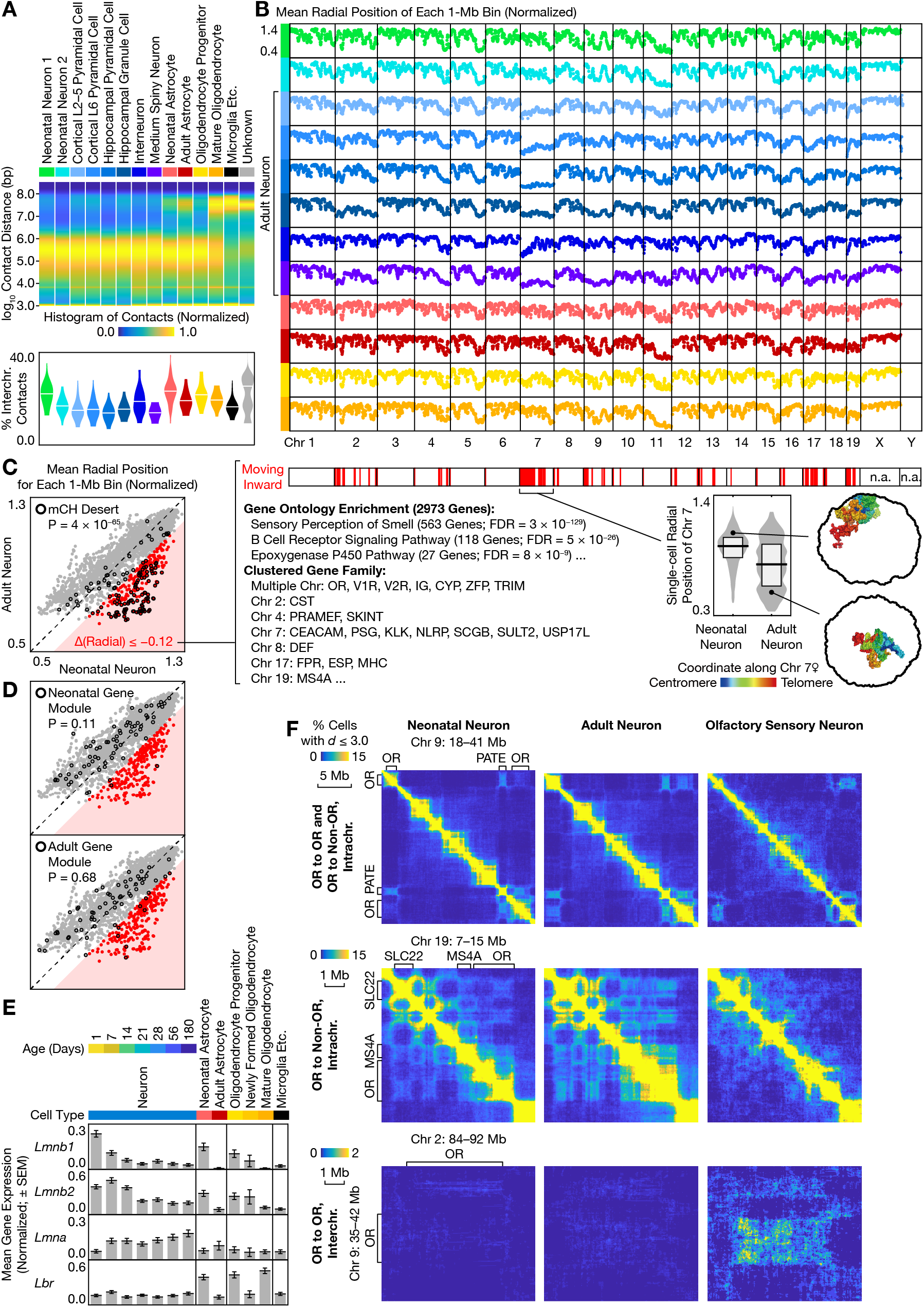
Large-scale 3D genome reorganization during postnatal development. (A) Histogram of contact distances (log10 values) (top), and distribution of percent inter-chromosomal contacts among single cells (bottom), for each structure type. For histograms (top): each histogram was normalized to the highest value; the horizontal stripe around 4.0 (10 kb) was artifacts caused by structural variations. For inter-chromosomal contacts (bottom): horizontal lines denoted the mean; *n* was the same as in Figure 3D; bandwidth was 2%. (B) Mean radial position of each 1-Mb bin for each structure type. Radial positions (3D distances to the nuclear center) were normalized to the mean in each cell. *n* = 237, 110, 94, 33, 74, 86, 57, 19, 27, 19, 20, 41 cells × 2 alleles, from top to bottom. If a bin was missing in some cells, an average was still calculated if *n* ≥ 5 alleles. Note that values along sex chromosomes might not be directly comparable, because P1 animals were female while the rest were male. (C) Comparison of mean radial positions between neonatal and adult neurons (left), characteristics and genome-wide distribution of inward-moving regions (middle), and radial positions of Chr 7 among single neurons (right). For radial comparison (left): *n* = 347 and 363 cells × 2 alleles for neonatal and adult neurons, respectively; if a bin was missing in some cells, an average was still calculated if *n* ≥ 80% of total alleles; P-value was from two-sided Fisher’s exact test. For Chr 7 (right): horizontal lines and boxes denoted the median and quartiles, respectively; *n* was the same as above; bandwidth was 0.03. OR: olfactory receptor. V1R: vomeronasal 1 receptor. V2R: vomeronasal 2 receptor. IG: immunoglobulin. CYP: cytochrome P450. ZFP: zinc finger protein. TRIM: tripartite motif-containing. CST: cystatin. PRAMEF: PRAME like. SKINT: selection and upkeep of intraepithelial T cells. CEACAM: carcinoembryonic antigen-related cell adhesion molecule. PSG: pregnancy specific glycoprotein. KLK: kallikrein. NLRP: NLR family, pyrin domain containing. SCGB: secretoglobin. SULT2: sulfotransferase family 2. USP17L: ubiquitin specific peptidase 17-like. DEF: defensin. FPR: formyl peptide receptor. ESP: exocrine gland secreted peptide. MHC: major histocompatibility complex. MS4A: membrane-spanning 4-domains, subfamily A. (D) Similar to (C) (left) but with the 2 correlated gene modules rather than mCH deserts. (E) Mean expression of lamin-related genes for each cell type. Expression levels were normalized before averaging (which therefore represented an intermediate between arithmetic and geometric means). *n* = 385, 343, 284, 362, 325, 268, 310, 230, 116, 177, 39, 314, 364 from left to right. (F) Example 3D proximity maps around clustered gene families. *n* = 347, 363, 106 cells × 2 alleles (for intra-chromosomal) or 4 allele pairs (for inter-chromosomal), from left to right. In the upper right panel, the bright yellow spot within the long-range OR-OR interaction corresponded to OR enhancers. PATE: prostate and testis expressed. SLC22: solute carrier family 22. Note that some regions were unmappable, leading to poor estimation of 3D distances.

In neurons, the distribution of contact distances stayed relatively unchanged during development (Figure 4A top), consistent with a previous bulk Hi-C study where chromatin domain sizes only changed by a small amount (777 kb in neural progenitors vs. 835 kb in neurons) during human neuron differentiation *in vitro* (Rajarajan et al., 2018). We further found the youngest structure type (neonatal neuron 1, mostly consisting of P1 cells) to harbor more inter-chromosomal contacts (Figure 4A bottom); this increase was uniform across the mouse genome (Figure S10). However, the trend became more complex when different ages and tissues were analyzed separately and when compared to additional data, suggesting possible batch effects (see later section) (Figure S16).

### Neurons undergo large-scale radial reconfiguration

We next looked at the most prominent difference between neonatal and adult neurons: radial positioning—preference for the nuclear periphery or interior—along the genome (Figure 4B). Dip-C is uniquely capable of measuring single-cell radial profiles, because other single-cell 3C/Hi-C methods cannot produce 3D structures, while the new GPSeq method (Girelli et al., 2020) has not been demonstrated in single cells.

We have previously used Dip-C to uncover unusual radial configuration in mouse sensory neurons—genome-wide inversion of euchromatin and heterochromatin in rod photoreceptors, and inward movement of olfactory receptor (OR) genes in olfactory sensory neurons (OSNs) (Tan et al., 2019)—both of which yielded highly consistent results with DNA *in situ* hybridization data (Clowney et al., 2012; Solovei et al., 2009).

In the brain, although the 6 adult structure types each exhibited a cell-type-specific radial profile, we found them to differ consistently from the neonatal types (Figure 4B). Many regions across the genome—including the majority of mouse Chr 7, 17, and 19—were relocated significantly inward from neonates to adults (Figure 4C, Table S10). For example, Figure 4C showed the distribution of Chr 7’s radial positions among neonatal and adult neurons. Such radial reorganization was absent from astrocytes and oligodendrocytes (Figure 4B bottom), and was therefore highly neuron-specific.

### Comparative analysis reveals convergent and divergent mechanisms behind neuron-specific 3D reorganization of clustered gene families

We suggested potential mechanisms for the neuron-specific radial reorganization by linking it to 2 known phenomena. First, regions that moved inward were highly enriched in previously identified “methylated CpH (mCH) deserts” (Lister et al., 2013) (Figure 4C left). To quantify this, we first identified the top 261 1-Mb bins (11% out of 2,391 autosomal bins examined) that moved inward during postnatal development (normalized radial position decreased by at least 0.12) (Table S10). Published mCH deserts corresponded to 129 bins, 92 (71%) of which were among the top inward-moving bins—a highly significant enrichment (71 vs. 11%; two-tailed Fisher’s exact test: P = 3.6 × 10^−65^).

These mCH deserts are low in CpG frequency and chromatin accessibility, and sometimes harbor large gene family clusters such as ORs, vomeronasal receptors (VRs, including V1Rs and V2Rs), and immunoglobulins (IGs) (Lister et al., 2013). Consistent with this, GO analysis of the 3.0 k genes in the top inward-moving regions revealed enrichment for OR- and IG-related terms (sensory perception of smell: 563 genes, FDR = 3 × 10^−129^; B cell receptor signaling pathway: 118, 5 × 10^−26^) (Figure 4C middle, Table S11). Further examination of inward-moving genes identified many other clustered gene families, such as cytochromes P450 (CYPs), zinc finger proteins (ZFPs), the major histocompatibility complex (MHC), and the membrane-spanning 4-domains subfamily A (MS4A) (Figure 4C middle, Table S11).

More importantly, mCH deserts are known to escape the neuron-specific, global non-CpG DNA methylation—a major epigenomic event during the same developmental window (P7–28) (Lister et al., 2013). This highlighted an intriguing relationship between DNA methylation and 3D genome structure, because regions that moved radially were primarily those left untouched by *de novo* DNA methylation (Figure 4C left).

Second, the specific radial movement of CpG-poor regions (Figure S11)—including OR and VR gene clusters—was reminiscent of our previous observation in OSNs. During OSN development, ORs move inward, detach from the nuclear lamin, and form both long-range and inter-chromosomal interaction to ensure their “one-neuron-one-receptor” expression (Clowney et al., 2012; Tan et al., 2019). Indeed, radial profiles were similar between adult neurons and OSNs (Figure S11). Adult neurons also harbored long-range interaction between OR gene clusters (Figure 4F top). Such long-range interaction was independently confirmed by a recent preprint using an orthogonal method (immunoGAM) (Winick-Ng et al., 2020).

Despite the striking similarity, our data also highlighted important differences between neurons in the brain and OSNs. In OSNs, long-range interaction is highly specific to ORs. In the brain, however, adult neurons additionally formed interaction between ORs and other clustered gene families, and between other families, with strength similar to (or even higher than) OR-OR interaction (Figure 4F top two rows). Furthermore, unlike OSNs, in the brain contacts were not sharply concentrated at OR enhancers (Figure 4F top). Finally, inter-chromosomal OR interaction was weaker in the brain than in the nose (Figure 4F bottom). In contrast, inter-chromosomal interaction between certain inward-moving regions—such as between VRs on Chr 7 and 17—was specifically strengthened in adult neurons to a much greater extent (Figure S12).

Such difference might be explained by divergent expression patterns of chromatin-regulating genes. In both OSNs and rod photoreceptors, developmental down-regulation of *Lbr* (lamin B receptor) plays a critical role in 3D genome transformation (Clowney et al., 2012). In our MALBAC-DT data, however, *Lbr* expression was low yet constant in neurons, only dynamic in astrocytes and oligodendrocytes (Figure 4E). In addition, *Lhx2*—whose expression is crucial for both long-range and inter-chromosomal OR interaction in OSNs (Monahan et al., 2019)—was sharply down-regulated around P14, as a member of the neonatal gene module (Figure 2F). Therefore, a similar yet distinct pathway must underlie our observed 3D genome reorganization. Interestingly, we found that both *Lmnb1/2* (lamin B1/2) were developmentally down-regulated in all 3 cell lineages, and *Lmna* (lamin A/C) somewhat up-regulated in neurons. These genes might contribute to the structural transformation.

Paradoxically, we found little correlation between gene expression changes and radial reorganization. Neither the neonatal nor the adult gene module (Figure 2F, Table S5) was enriched in inward-moving regions (two-sided Fisher’s exact test: P = 0.11 and 0.68, respectively) (Figure 4D). However, 3D reorganization of ORs might help further suppress OR expression, which is prevalent in embryonic stem cells (ESCs) yet rare in the brain, and might be related to OR expression in a small, activated population of hippocampal pyramidal cells, as a recent preprint suggested (Winick-Ng et al., 2020).

### Developmental chromatin A/B compartmentalization is correlated with gene expression modules

In addition to large-scale radial reconfiguration, the genome also refolded on finer scales during the 3D transformation. To investigate these changes, we began by identifying genomic regions with developmentally regulated scA/B values, which together underlay the separation of neonatal and adult structure types on the t-SNE plot (Figure 3A). In glia, we have already shown (see earlier section) that scA/B differences correlated well with gene expression. We now focused on neurons.

In neurons, we identified the top 100 1-Mb bins (out of 2,410 autosomal bins examined) with developmentally down- and up-regulated scA/B values, respectively, by comparing adult structure types as a whole to neonatal ones (Figure 5A left). These regions were distributed across the mouse genome (Figure 5B, Table S8), and had very few enriched GO terms (Table S11). The 869 genes with decreasing scA/B (becoming more B-like and more heterochromatic during development) were enriched in sensory perception of smell (driven by OR clusters on multiple chromosomes; 188 genes; FDR = 8 × 10^−56^); other enrichments were driven by individual gene clusters, and may therefore be spurious (Table S11).

**Figure 5.**
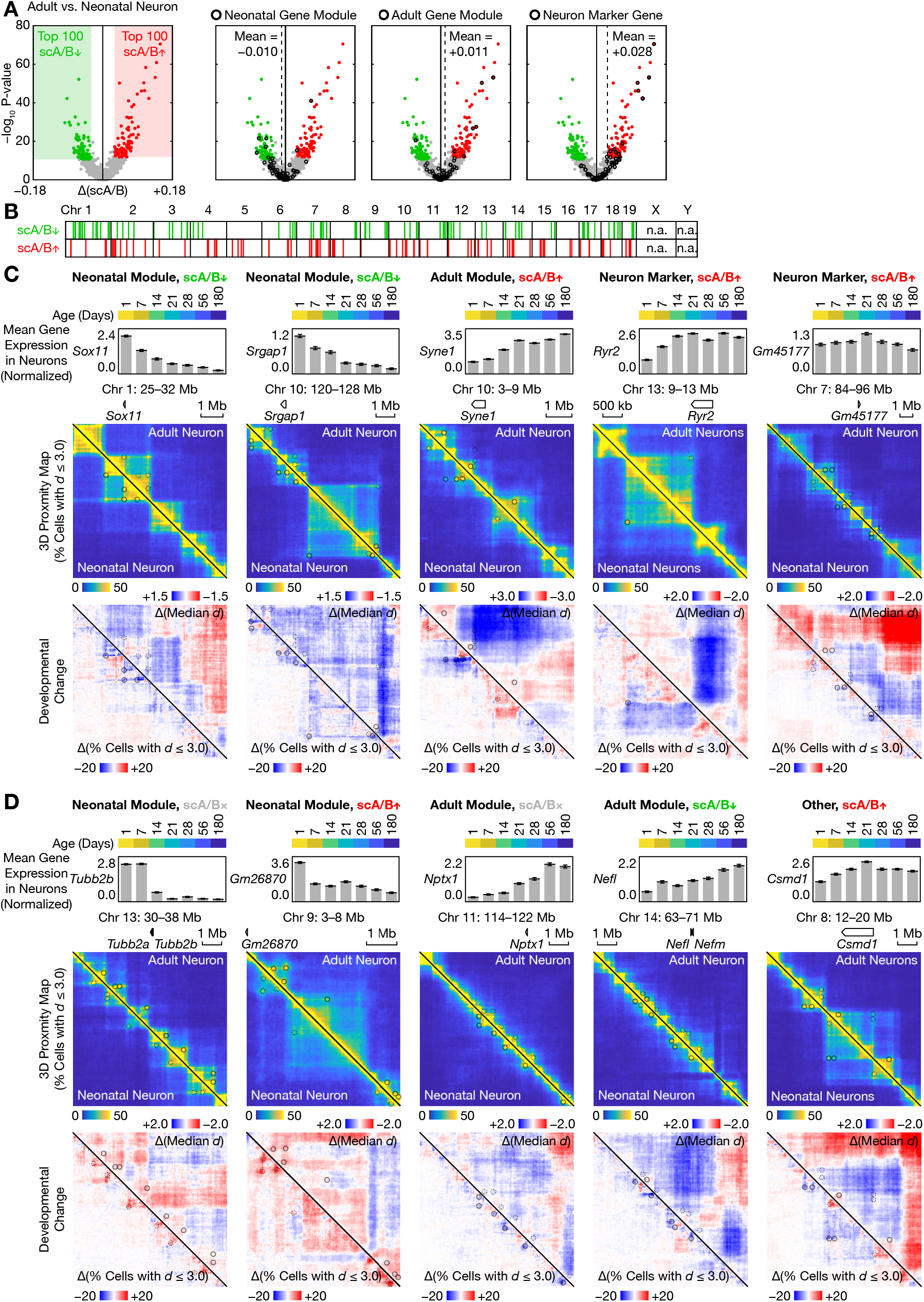
Correlation between developmental changes in 3D genome and transcriptome. (A) Volcano plot of differential scA/B between neonatal and adult neurons. P-values were from two-sided Mann–Whitney U tests. *n* = 795 vs. 585. Top 100 regions on each side were selected from regions with a minimal scA/B difference of 0.03, sorted by P-values. (B) Genome-wide distribution of differential scA/B regions. (C) Mean expression among neurons at each age (top), 3D proximity maps (middle), and developmental structural changes (bottom) for example genes with concordant changes in scA/B and expression. Circles denoted changing chromatin loops. For mean expression (top), error bars denoted SEM; *n* = 385, 343, 284, 362, 325, 268, 310 from left to right. For other panels (middle and bottom), *n* = 363 vs. 347 cells × 2 alleles. Note that some regions were unmappable, leading to poor estimation of 3D distances. Also note that *Ryr2* (second to right) was classified as part of the adult module under less stringent criteria (Table S6), and that *Gm45177* (right) resided in an intron of *Dlg2* and might represent a neuron-specific isoform of *Dlg2* rather than a separate non-coding gene. (D) Similar to (C), but for example genes with discordant changes in scA/B and expression.

We then compared scA/B changes to gene expression. On average, the two developmentally regulated gene modules—the neonatal and adult modules—changed scA/B by −0.010 and +0.011, respectively, consistent with the directions of their expression changes (Figure 5A middle). The less stringent gene list (Table S6) yielded similar results (scA/B changed by −0.009 and + 0.009). For individual 1-Mb bins, scA/B values increased (regardless of statistical significance) for 38% (26 out of 68) and 61% (39 out of 64) of bins containing neonatal and adult genes, respectively. Both represented significant differences from the whole genome (38 vs. 50%, and 61 vs. 50%; two-sided Fisher’s exact test: P = 7.3 × 10^−9^ for both) that were consistent with the directions of their expression changes. Taken together, we found correlated regulation of A/B compartments and gene expression during postnatal neuron development.

Interestingly, neuron-specific marker genes—the majority of which were already highly expressed at birth—presented a better match to developmental scA/B changes than the adult gene module—genes that were actually up-regulated (Figure 5A right). On average, neuron marker genes changed scA/B by +0.028 during postnatal development, much higher than +0.011 for the adult module. These genes also corresponded to a higher fraction of individual bins (75%; 60 out of 80) that increased in scA/B. This suggested that 3D structure of cell-type-specific genes might undergo further remodeling after their initial up-regulation during cell-type commitment.

We visualized local 3D genome rewiring around example genes with either concordant (Figure 5C) or discordant (Figure 5D) changes in scA/B and expression. In particular, we quantified spatial reorganization by both changes in short-distance interactions—defined as the fraction of cells where each pair of 20-kb particles are nearby in 3D (distance *d* ≤ 3.0 radii), which we termed the “3D proximity map” (equivalent to contact maps in bulk Hi-C, but with an absolute scale and clear 3D interpretation) (Tan et al., 2019), and changes in longer-distance interactions—defined as the median 3D distance (which cannot be obtained with other single-cell 3C/Hi-C methods). This allowed us to visualize complex 3D changes such as separation of adjacent domains and changing enhancer-promoter loops. Other examples of prominent local 3D rewiring were shown in Figure S13.

### Sensory deprivation does not affect 3D genome transformation

Our striking observation of postnatal 3D genome transformation leads to several questions: (1) Given the intriguing timing (between P7 and 28)—a critical period during which the brain is highly plastic and influenced by myriad sensory inputs (for example, eyes open around P12), is the 3D genome transformation dependent on sensory experience, or genetically predetermined? (2) Is the transformation continuous, or abrupt at some time point, between P7 and 28? (3) How robust are our findings against experimental variables?

To answer these questions, we performed Dip-C on 1,692 single cells from the visual cortex of dark-reared and control mice at 5 different ages in the first postnatal month: on P1, P7, P14, P21, and P28 (Figure 1B bottom). Similar to the main dataset, we obtained an average of 323 k chromatin contacts per cell (SD = 93 k, minimum = 101 k, maximum = 963 k) (Figure S14, Table S7). This design allowed us to determine the experience dependence of 3D genome transformation by comparing dark-reared (which dramatically reduced sensory inputs to the visual cortex) and control mice, and to analyze fine temporal dynamics with weekly resolution. Note that at the earliest time point (P1, the day when dark rearing started), only control mice were studied.

To test the robustness of our results, we changed both the mouse strain (from CAST × B6 F1 hybrids in the main dataset, to the widely used B6 inbred strain) and the restriction enzyme in Dip-C (from GATC-cutting MboI in the main dataset, to CATG-cutting NlaIII, which was recently used by another single-cell 3C/Hi-C method snm3C-seq (Lee et al., 2019)) (Methods).

Despite complete elimination of visual inputs, 3D genome transformation in all 3 cell lineages proceeded normally in the visual cortex. On the t-SNE plot, cells formed all the normal structure type clusters, with dark-reared and control cells relatively intermixed in each (Figure 6A). Among glia, adult structure types emerged on P14 in both experimental groups, albeit with variable fractions (Figure 6B). Among neurons, both experimental groups transitioned normally from neonatal structure types to the same 3 adult types on P28—cortical pyramidal cells, interneurons, and MSNs (inferred via joint t-SNE with the main dataset (Figure S15) and scA/B analysis).

**Figure 6.**
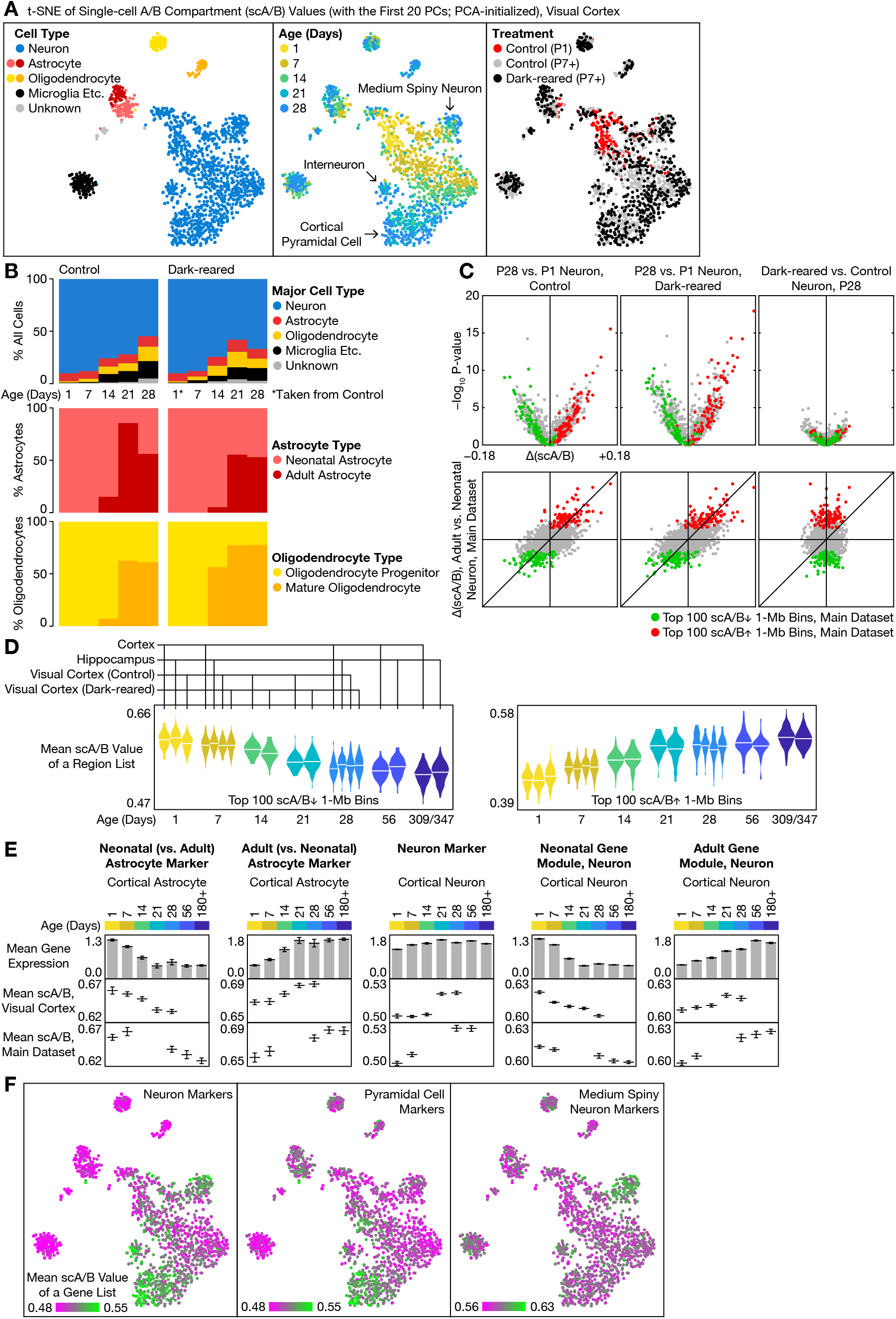
Single-cell 3D genome in the visual cortex of dark-reared and control mice. (A) t-SNE of single-cell 3D genomes. t-SNE was performed on the first 20 PCs of scA/B values. (B) Composition of structure types for each tissue and age. For all cells (top), *n* was shown in Figure 1B. For astrocytes (middle): *n* = 15, 14, 14, 16, 18, 15, 10, 16, 22, 18 from left to right. For oligodendrocytes (bottom): *n* = 2, 6, 13, 14, 25, 2, 7, 18, 27, 17. (C) Volcano plots for differential scA/B between ages or treatments. P-values were from two-sided Mann–Whitney U tests. *n* = 100 vs. 171 (left), 127 vs. 171 (middle), 127 vs. 100 (right). (D) Distribution of mean scA/B values of top regions with developmentally regulated scA/B, among single neurons for each tissue and age. Horizontal lines denoted the mean. *n* = 157, 178, 171, 125, 120, 169, 169, 140, 141, 135, 108, 89, 55, 100, 127, 111, 111, 225, 209 from left to right. Bandwidth was 0.005. (E) Mean expression (top) and mean scA/B values (bottom two rows) of example gene lists, among single cells at each age. Error bars denoted SEM. For cortical astrocytes (left 2 columns), *n* = 38, 31, 23, 13, 14, 22, 6, 15, 24, 30, 38, 36, 14, 11, 29, 17, 14 from left to right and from top to bottom. For cortical neurons (right 3 columns), *n* = 205, 196, 83, 184, 169, 151, 171, 171, 338, 281, 243, 227, 157, 125, 89, 111, 225. (F) Mean scA/B values of example gene lists, among single cells on the t-SNE plot.

We found little effect of sensory deprivation on 3D genome, even with detailed analysis of scA/B differences in neurons (Figure 6C). At each age examined, dark rearing led to little change in scA/B (Figure 6C right). Out of 2,397 autosomal bins examined, only 3, 2, 6, and 4 bins reached a threshold of FDR ≤ 0.05 (or 2, 1, 3, and 0 bins at FDR ≤ 0.01; two-sided Mann–Whitney U tests with Benjamini–Hochberg procedure) on P7, 14, 21, and 28, respectively. None of the differential bins overlapped between ages. To rule out insufficient statistical power, we contrasted the above result by comparing P1 and 28 within each experimental group (Figure 6C left two columns), where 550 and 797 bins reached FDR ≤ 0.05 in control and dark-reared mice, respectively.

Developmental changes in scA/B were highly consistent with our main dataset. In neurons, scA/B differences between P1 and 28 in both experimental groups were highly correlated with those between neonatal and adult neurons in our main dataset (Pearson’s r = +0.61 and +0.59 in control and darked-reared mice, respectively) (Figure 6C bottom). Correlation was even higher (r = +0.65) when tissue (cortex) and ages (P1 and 28) were matched in the main dataset. Therefore, despite a small batch effect (as evidenced by a higher r = +0.84 between the two 2 experimental groups in the new dataset; presumably caused by different mouse strains, restriction enzymes, and cortical regions), all our major findings remained unchanged in the new dataset. This suggested that 3D genome transformation was genetically predetermined, unaffected by early-life experience.

### 3D genome transformation is continuous in the first postnatal month

The high temporal resolution of our new dataset provided a detailed look into the dynamics of 3D genome transformation. In neurons, our main dataset only captured the 2 discontinuous end points of their prominent structure type conversion. The intermediate steps, however, remained unclear. Does the conversion involve many continuously evolving, intermediate structure types? Or is it a discrete jump between the 2 end states?

Distribution of cells on the new t-SNE plot suggested gradual—rather than abrupt—3D genome transformation during the first month of life (Figure 6A). In particular, neurons formed a continuous trajectory, with a clear age gradient from P1 to 28, on the t-SNE plot (Figure 6A middle). We further quantified this process by calculating for each age the mean scA/B values of the top 100 bins with developmentally up- and down-regulated scA/B, respectively, from the main dataset. In our new dataset, mean scA/B values of both categories (up and down) changed continuously (and regardless of dark rearing), with end points matching the main dataset (Figure 6D). Note that scA/B changes continued after P28, albeit with a much slower rate (Figure 6D).

The new t-SNE plot also confirmed earlier structural specialization of certain sub-types. In our main dataset, some granule cells could be seen relatively clustered on P1 and 7 (see earlier section). With better temporal continuity of our new dataset (which would facilitate cell clustering and t-SNE by harboring more mutually similar cells), MSNs could also be seen splitting from the main structure type (mostly cortical pyramidal cells) as early as P1 (Figure 6A middle).

Our new dataset also enabled side-by-side temporal analysis of gene expression and scA/B. In astrocytes, for example, mean scA/B values of neonatal and adult markers faithfully tracked gene expression dynamics, changing gradually around P14 (Figure 6E left two columns). Mean scA/B values for various categories of neuronal genes (Figure 6E, Figure 6F) also correlated with gene expression changes, albeit with variable temporal concordance.

Finally, we examined the percentage of inter-chromosomal contacts (which reflected the overall extent of chromosomal intermingling) for neurons from each tissue and age in both datasets (Figure S16). In all experiments (cortex, hippocampus, visual cortex), the inter-chromosomal percentage was the highest at a neonatal age (P1 or 7), and the lowest at an intermediate age (P14 or 28). However, significant quantitative variation was observed, suggesting possible batch effects.

### Many imprinted loci adopt allele-specific 3D structures, one of which is particularly long-range

Allele-specific gene expression plays a crucial role in the brain and throughout mammalian development. We have previously used Dip-C to study random monoallelic expression of female X chromosomes (i.e. X chromosome inactivation (XCI)) in a human B-lymphoblastoid cell line, via haplotype-resolved scA/B analysis (Tan et al., 2018). Here we similarly analyzed female cells (P1 animals in the main dataset) and observed two imbalanced (265 vs. 139 cells; 66 vs. 34%) clusters along the first PC of haplotype-resolved Chr X scA/B values (Figure S17), consistent with known biased XCI in F1 hybrids (more cells choose the CAST allele than the B6 one).

Genome imprinting presents another important example of allele-specific expression, which suppresses a specific parental allele of each imprinted gene; however, its molecular mechanism is not fully understood. Scattered evidence has suggested parent-of-origin-specific 3D genome structure as a factor. For example, we have previously visualized allele-specific chromatin loops at an imprinted locus (*H19*/*IGF2*) in the human blood (Rao et al., 2014; Tan et al., 2018). However, allele-specific 3D genome structure has not been systematically examined across the genome.

Here we studied parent-of-origin-specific 3D genome structure around all known imprinted genes (in an 8-Mb region around each gene or gene cluster), by comparing reciprocal crosses (two crosses with parental strains swapped). Reciprocal crosses can eliminate all confounding factors, because only true parent-of-origin effects will show consistent parental differences between the two crosses (CAST/EiJ♀ × C57BL/6J♂ and C57BL/6J♀ × CAST/EiJ♂), while strain-specific confounders (primarily artifacts caused by strain-specific copy-number/structural variations, but also potentially strain-specific 3D structures) will show opposite signs (Figure 7A).

**Figure 7.**
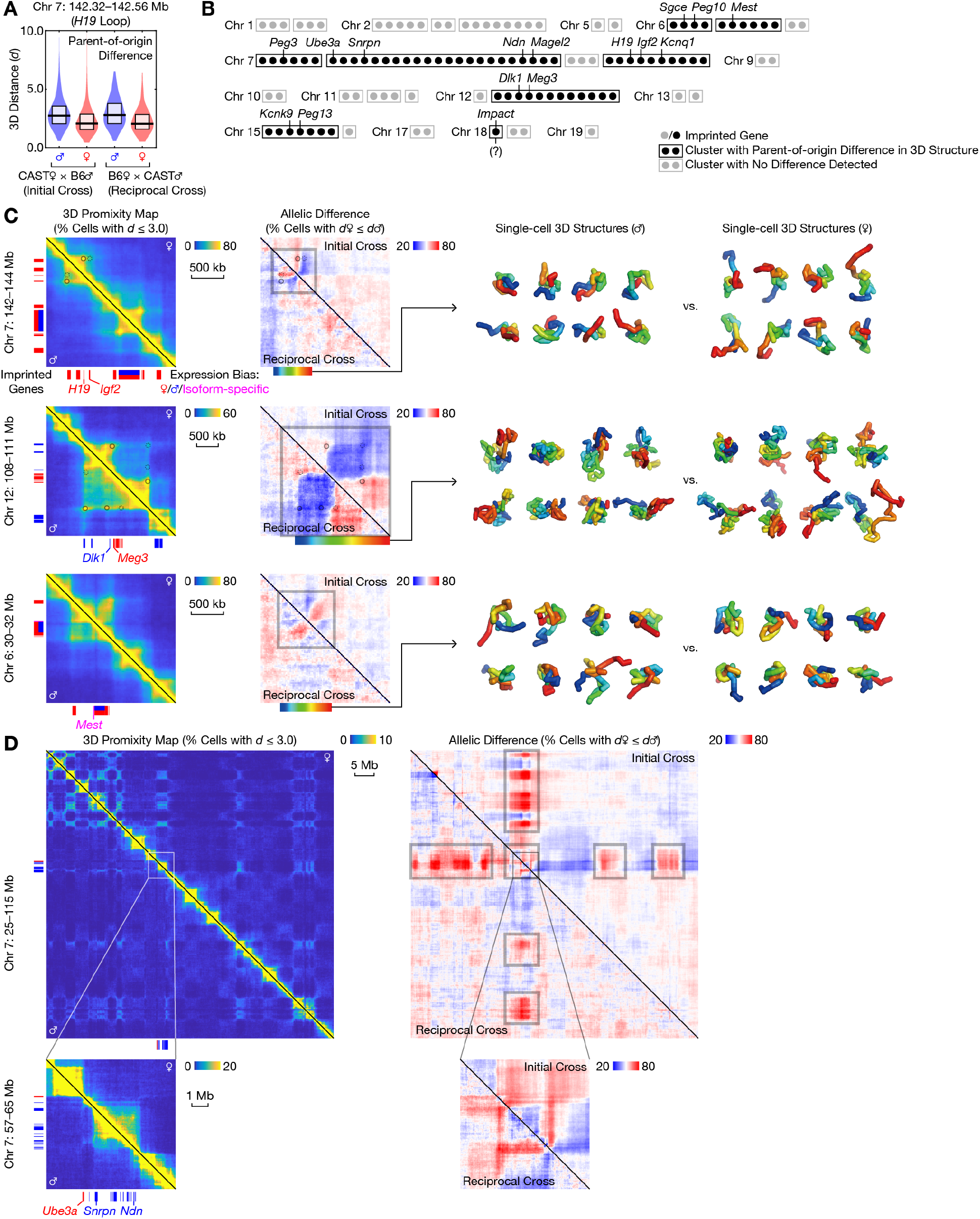
Parent-of-origin-specific 3D genome structure in imprinted gene clusters. (A) Distribution of allele-specific 3D distances for an example parent-of-origin-specific chromatin loop (at the *H19*/*Igf2* locus). Horizontal lines and boxes denoted the median and quartiles, respectively. *n* = 666 and 168 for the initial and reciprocal crosses, respectively. (B) Summary of detected parent-of-origin differences among imprinted gene clusters. (C) 3D proximity maps (left), allelic differences (middle), and example 3D structures (right) of example imprinted gene clusters with parent-of-origin-specific 3D structure. For 3D proximity maps (left), *n* = 834. For allelic differences (middle), *n* was the same as in (A). Gray boxes denoted regions with clear parent-of-origin-specific structures. Rainbows denoted regions to be visualized on the right (cells were selected randomly). (D) Similar to (C) but for the PWS/AS locus. Note that some regions (for example, between *Ube3a* and *Snrpn*) were unmappable, leading to poor estimation of 3D distances.

Out of the 29 clusters of known imprinted genes in the mouse brain (115 genes) (Perez et al., 2015), we found strong evidence for parent-of-origin-specific 3D genome structure in 7 clusters and weak evidence in 1 (Figure 7B). We quantified allele-specific differences with a non-parametric statistic—the fraction of cells where the paternal 3D distance was greater than the maternal one (P(*d*♀ ≤ *d*♂))—for each pair of 20-kb particles (Figure 7C, Figure S18, Figure S19). This statistic can only be measured by Dip-C but not other single-cell 3C/Hi-C methods, and has the unique advantage of only comparing 3D distances within a cell (not between cells)— thus canceling out any cell-type- or cell-state-specific confounders. In all 8 clusters, similar parent-of-origin difference existed in both neonatal (P1 and 7) and adult (P28 onward) cells (Figure S20). We further confirmed a previously reported, quantitative developmental difference (i.e. smaller parent-of-origin difference in the 3D size of the *Snrpn*–*Ube3a* region on P1) in a DNA *in situ* hybridization study of the Prader-Willi/Angelman syndrome (PWS/AS) locus (Leung et al., 2009). In comparison, no parent-of-origin difference was observed in 29 randomly selected 8-Mb regions across the mouse genome.

All our observed parent-of-origin effects were relatively local, with one notable exception: the PWS/AS locus on Chr 7 (corresponding to human Chr 15). Two alleles of this imprinted cluster not only exhibited parent-of-origin-specific domain organization within the ∼3-Mb cluster (Figure 7D bottom), but also interacted differently with multiple regions across the chromosome (Figure 7D top). In particular, only the maternal allele interacted with multiple Mb-sized, compartment B-like regions across Chr 7—up to tens of Mb away. Such difference was also apparent from raw contact maps, which did not involve any haplotype imputation or 3D modeling (Figure S21). This surprisingly long-range interaction provided new information about the molecular difference between the two alleles of this large imprinted gene cluster.

## DISCUSSION

Cells are the building blocks of the brain. According to the central dogma of molecular biology, a cell’s function is governed by its own “brain” —the 2 meters of genomic DNA folded in its nucleus, and the RNA molecules that it transcribes. However, previous studies could not resolve the 3D genome structures of single brain cells. This lack of knowledge presents a major hurdle for understanding the “structure-function” relationship of the mammalian genome, and for treating the many chromatin- and transcription-related developmental disorders. Our 3D genome and transcriptome atlases directly tackled this problem, by uncovering the 3D genome “structure type” behind each of the brain’s highly specialized cell types, tracing their developmental origins, and probing their relationship with dynamic gene expression and early-life experience.

Development is a highly dynamic process. Previous studies—single-cell or bulk—have not followed the temporal progression of cell-type-specific 3D genome structure in the brain *in vivo*. Moreover, the postnatal dynamics of single-cell transcriptome has not been systematically studied in a mammalian brain. By charting a year-long time course of 3D genome structure and its accompanying transcriptome progression, our database offered unique insights into the dynamic interplay between functional, anatomical, transcriptional, and structural cell types. Our discovery of postnatal transcriptome and 3D genome “differentiation” of neurons—rather late compared to their initial cell-type specialization—highlights an interesting possibility that 3D genome structure may act to reinforce and refine other forms of differentiation that precedes it. This developmental strategy can be especially useful for neurons, which generally do not self-renew and must perform highly specialized functions for life.

The timing of our observed 3D genome transformation is particularly intriguing. Mammals are born with a highly plastic brain that continues to develop its myriad cognitive functions over time. During this critical perinatal period, the brain undergoes numerous molecular and functional changes. For example, the majority of synaptogenesis, myelination, and perineuronal net formation occur during this developmental window. As the animal begins to receive various sensory inputs, its sensory processing (for example, in visual and auditory cortices) also exhibits unique “critical periods” of functional plasticity around this time. Inside the cell nucleus, the epigenome undergoes a neuron-specific episode of global non-CpG DNA methylation. Our discovery of a concurrent major event in transcriptome and 3D genome reorganization provides another molecular underpinning for the brain’s unusual cognitive plasticity during this time, offering new possibilities for shifting or extending this critical period. Our determination of its sensory independence further illustrates the molecular logic of brain development, where genetically predetermined molecular changes may drive the brain’s functional maturation.

Finally, our results provide potential therapeutic values for the treatment of developmental disorders. First, we uncovered both transcriptome and 3D genome maturation as new molecular distinction between fully functional, adult neurons and their neonatal, embryonic, and *in vitro* (stem-cell-derived) counterparts. This may offer a new direction for improving stem cell therapies for neurodevelopmental and neurodegenerative diseases. Second, our observed similarity between the brain and the nose in their 3D genome dynamics not only suggests candidate regulators for this shared process, but also enables further interrogation using the nose as a simple, well-characterized model system. The nose additionally offers easy access (via nasal biopsy) with minimal damage (neurons are regenerated throughout life) for clinical diagnosis and therapies. Lastly, our systematic survey of allele-specific genome structure suggests the involvement of 3D genome dysregulation in many imprinting disorders. Targeted correction of 3D genome abnormalities may therefore complement recent methylation-editing-based restoration of normal imprinting.

In summary, our discovery of cell-type-specific transcriptome and 3D genome transformation during a critical perinatal period provides a new molecular basis for neural plasticity, and opens up opportunities for 3D genome-based treatments for neurodevelopmental and genomic imprinting disorders.

## Supporting information

Supplementary Table Titles

## ACKNOWLEDGMENTS

This work was supported by Beijing Advanced Innovation Center for Genomics (ICG) at Peking University. L.T. was supported by a School of Medicine Dean’s Postdoctoral Fellowship and a Walter V. and Idun Berry Postdoctoral Fellowship from Stanford University. K.D. was supported by a grant from the Gatsby Foundation. The authors thank the Bauer Core Facility at Harvard University (Z. Niziolek, J. Nelson, and C. Reardon) for flow sorting and PCR machines, F. Alt, P. Wei, F. Gage, and S. Parylak for advice on nuclei isolation, and H. Li, X. Jin, Y. Chen, and L. Fan for helpful discussions.

## AUTHOR CONTRIBUTIONS

L.T., W.M., H.W., Y.Z., D.X., R.C., N.D., K.D., and X.S.X. designed the experiments. L.T., W.M., H.W., Y.Z., D.X., R.C., and N.D. performed the experiments. X.L. pre-processed MALBAC-DT data. L.T. analyzed the data. L.T., K.D., and X.S.X. wrote the manuscript.

