## Supplementary Figures for "Experience-independent transformation of single-cell 3D genome structure and transcriptome during postnatal development of the mammalian brain"

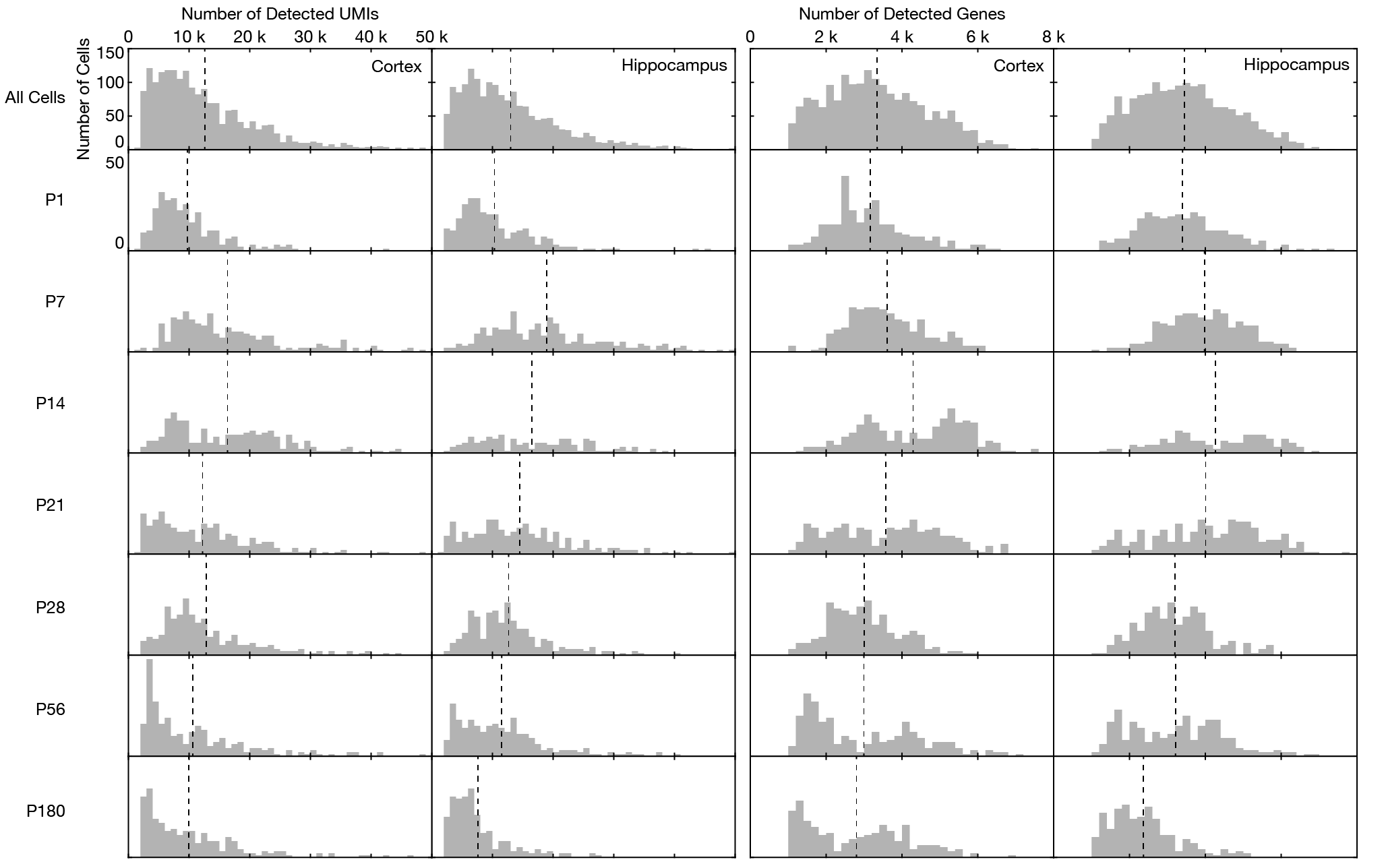

**Figure S1. Histograms of numbers of detected UMIs and genes per cell in the MALBAC-DT dataset.**

Bin size was 1 k for UMIs, and 0.2 k for genes. Dashed lines denoted the mean.

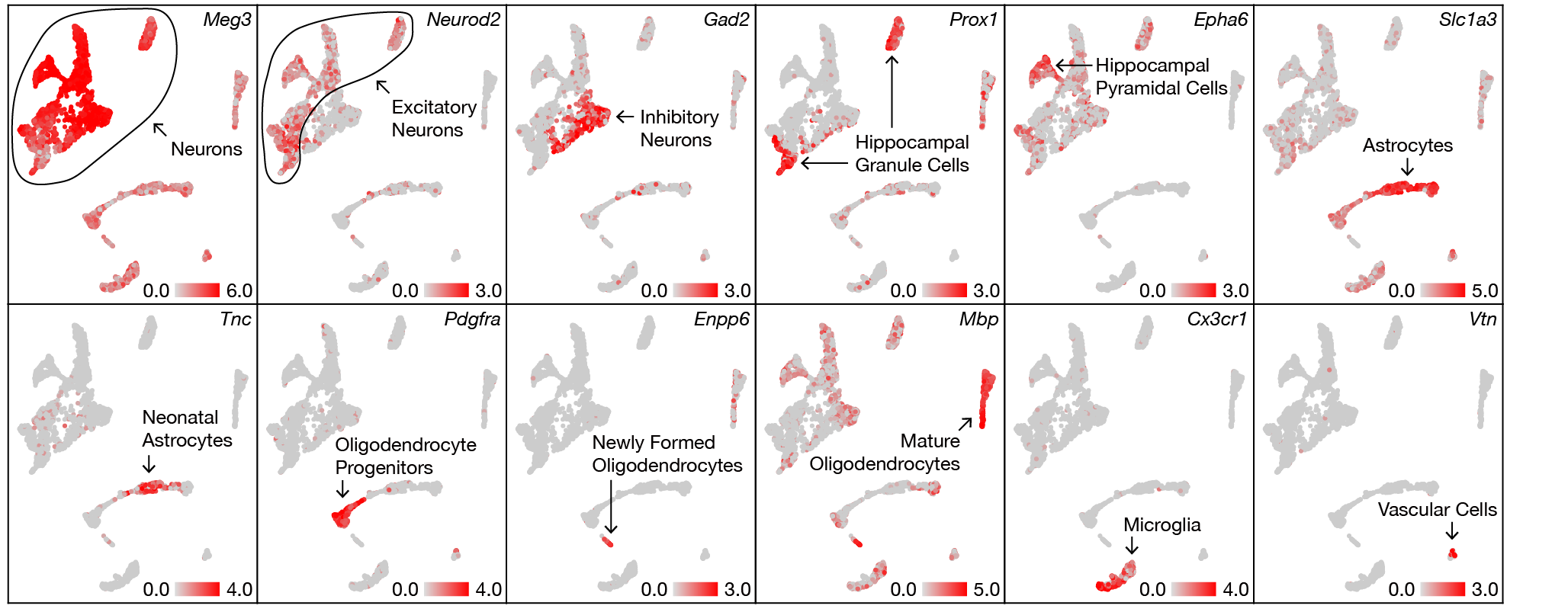

**Figure S2. Expression patterns of known marker genes on the UMAP plot of all cells.**

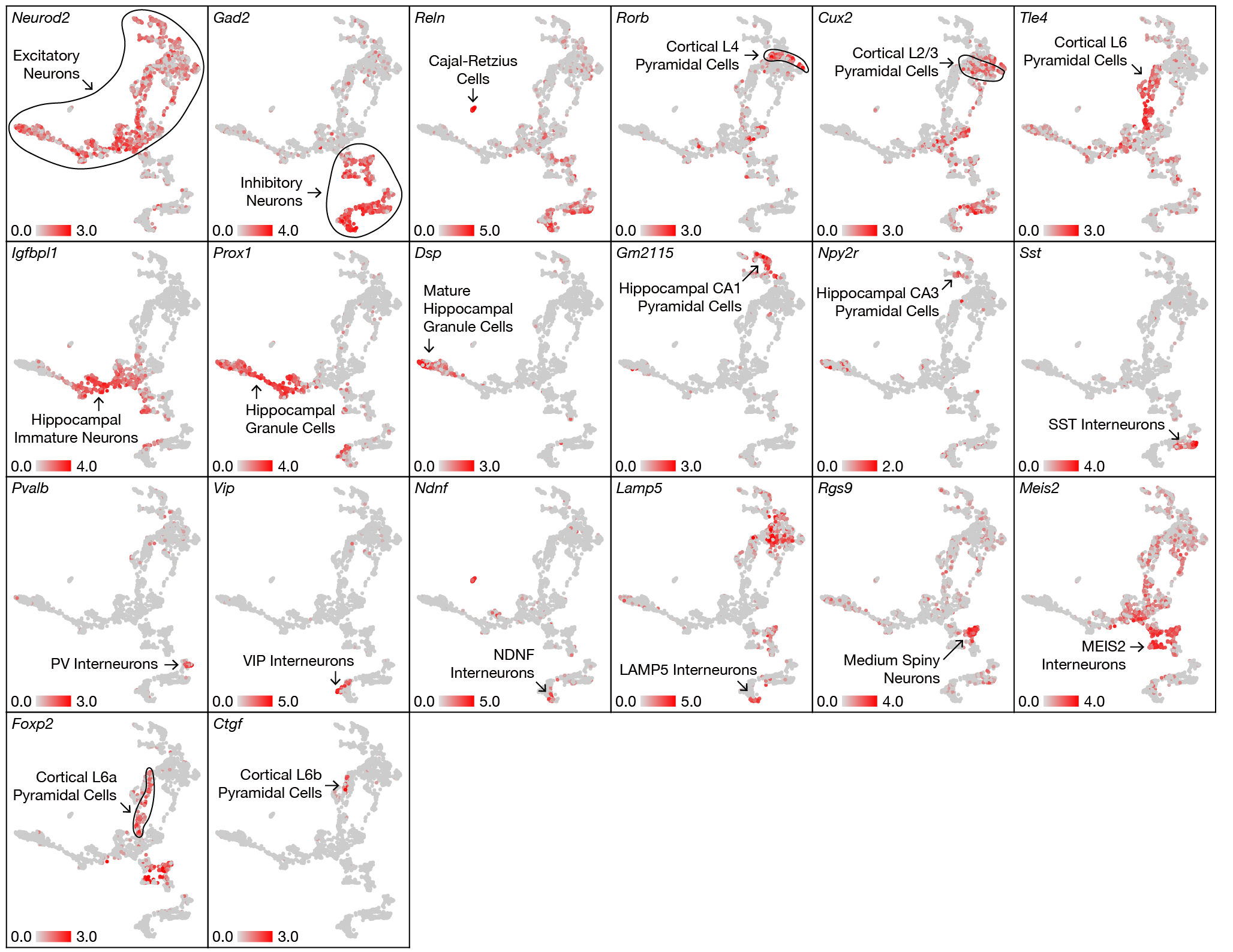

**Figure S3. Expression patterns of known marker genes on the UMAP plot of neurons.**

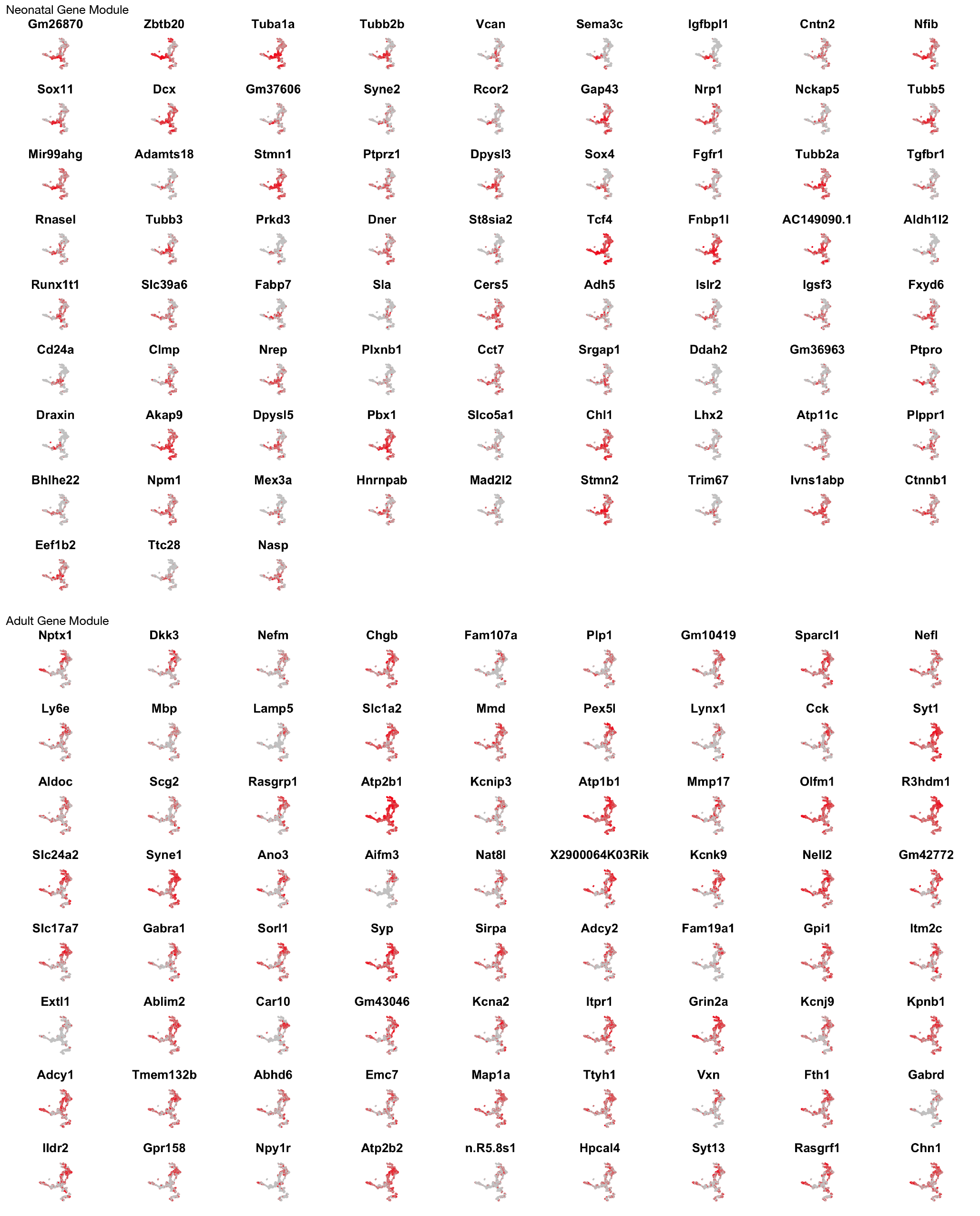

**Figure S4. Expression patterns of genes in the neonatal and adult modules, on the UMAP plot of neurons.**

Red denoted high expression. Color scale was set automatically by Seurat for each gene.

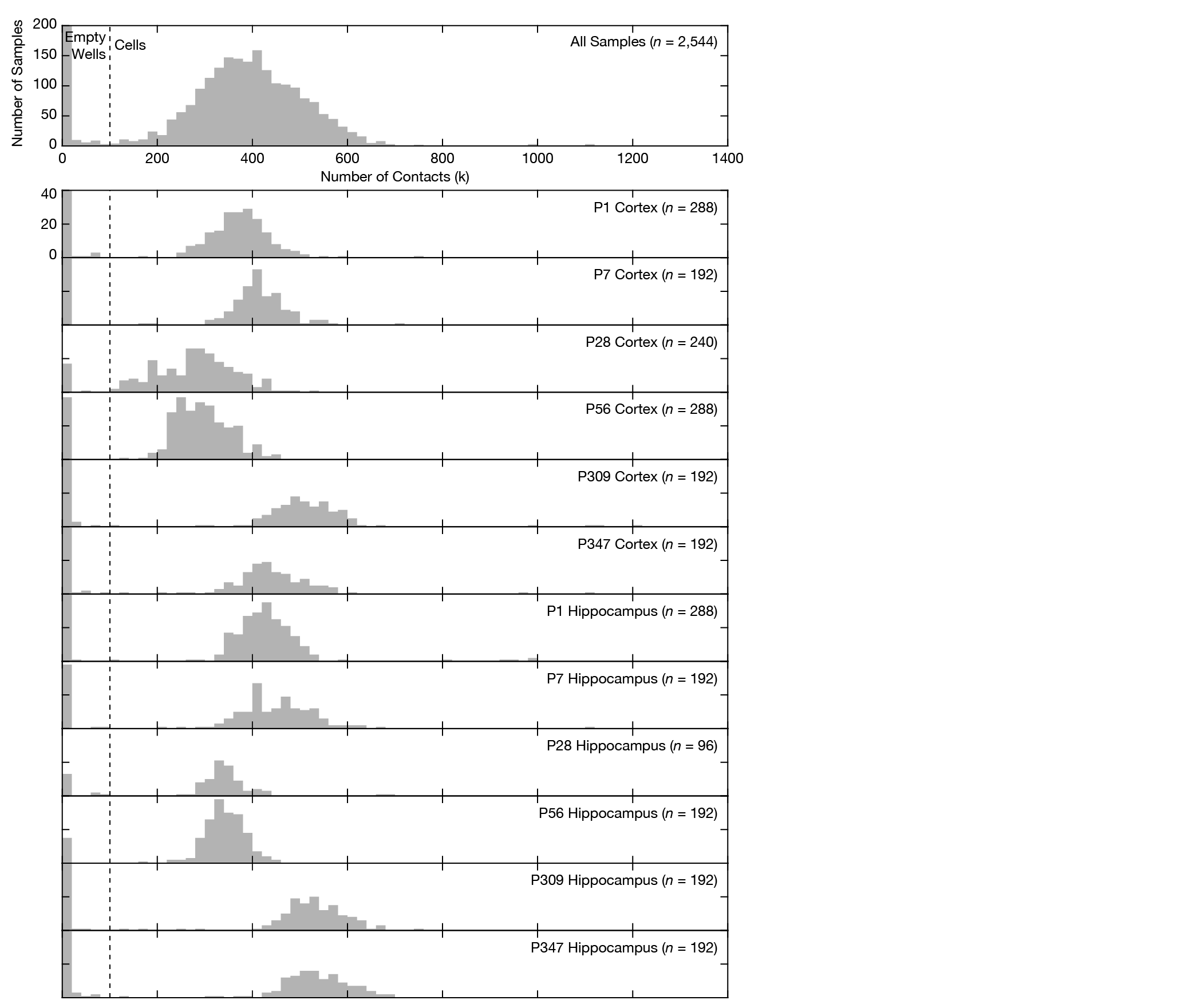

**Figure S5. Histograms of numbers of detected contacts per cell in the main Dip-C dataset.**

Bin size was 20 k. Dash lines denoted a threshold of 100 k, below which a sample was considered an empty well. The leftmost bins (0–20 k) were truncated in height for visual clarity.

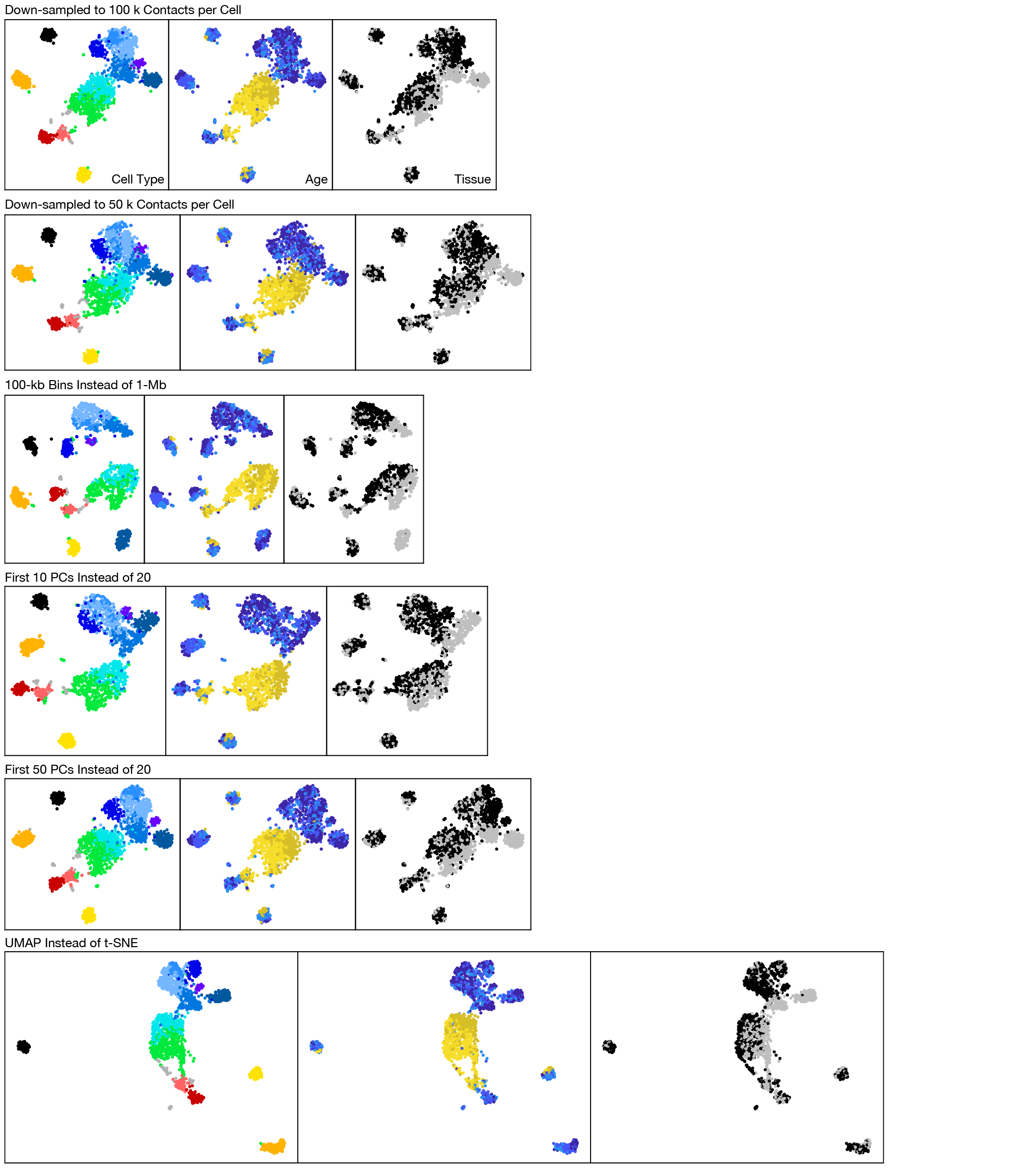

**Figure S6. Visualization of structure types with alternative choices of parameters.**

Coloring was the same as in Figure 3A.

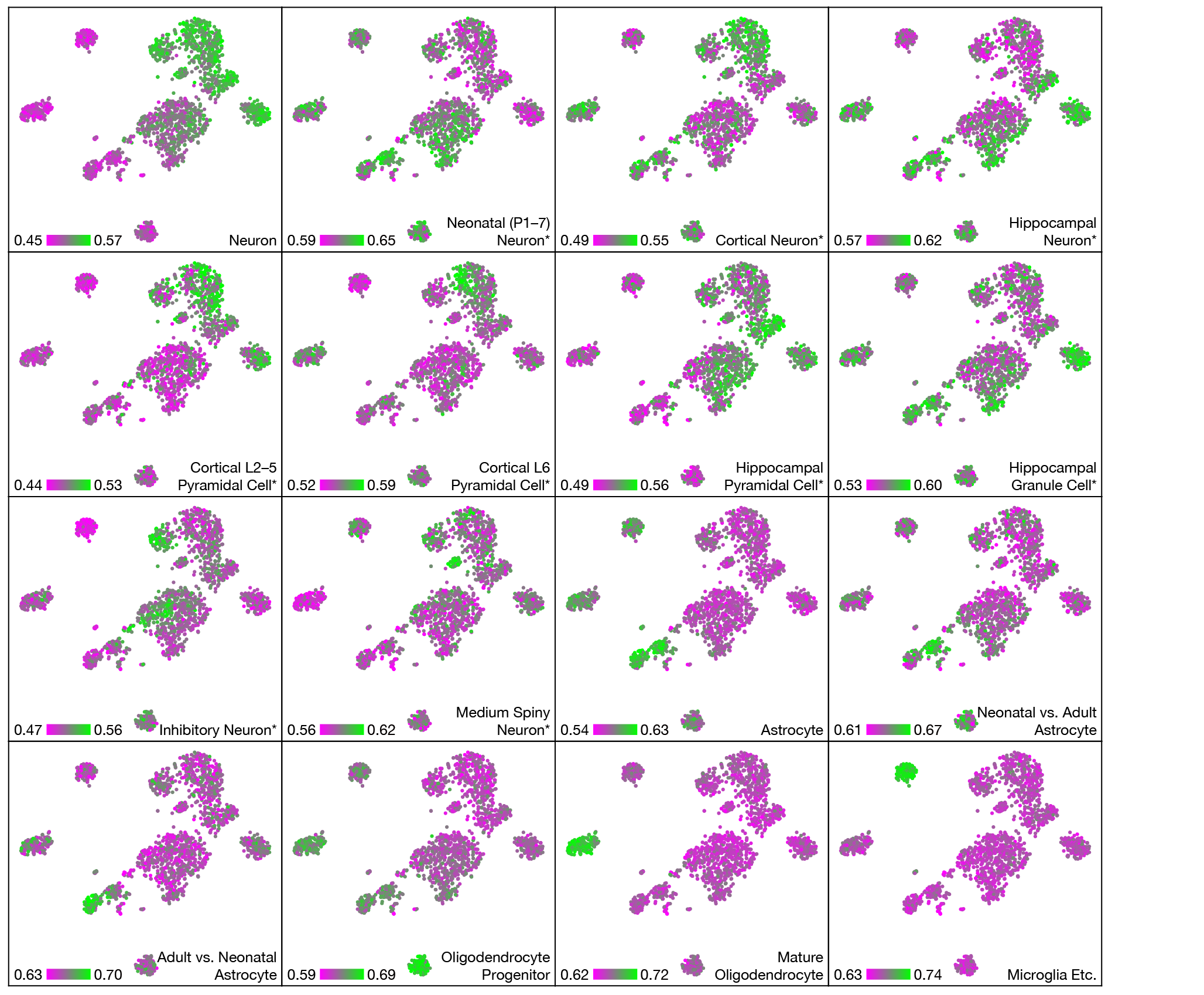

**Figure S7. Mean scA/B values of gene lists on the t-SNE plot.**

Similar to Figure 3D left, but plotted on the t-SNE plot of the main Dip-C dataset.

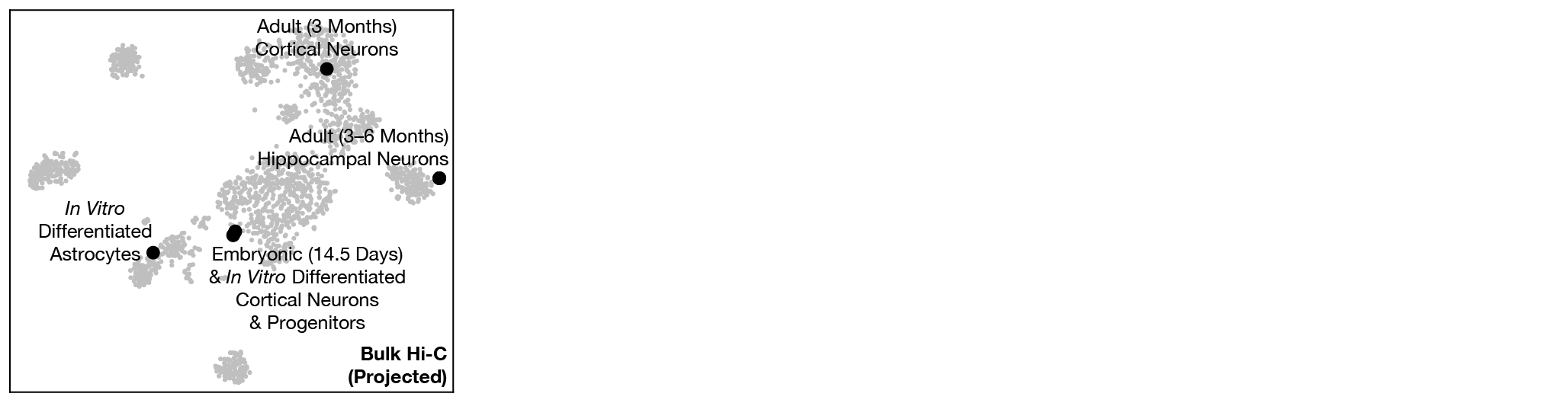

**Figure S8. Projection of published bulk Hi-C data onto the t-SNE plot.**

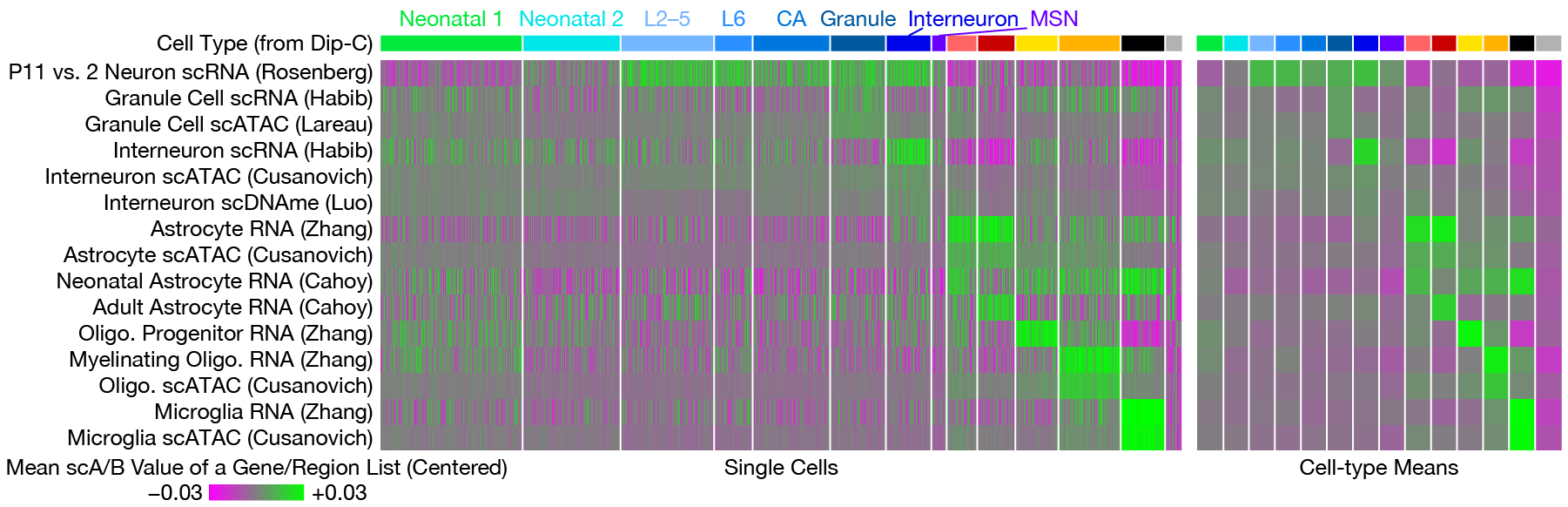

**Figure S9. scA/B analysis of gene/region lists from published omic data.**

Similar to Figure 3D left, but with published omic data. Oligo.: oligodendrocyte.

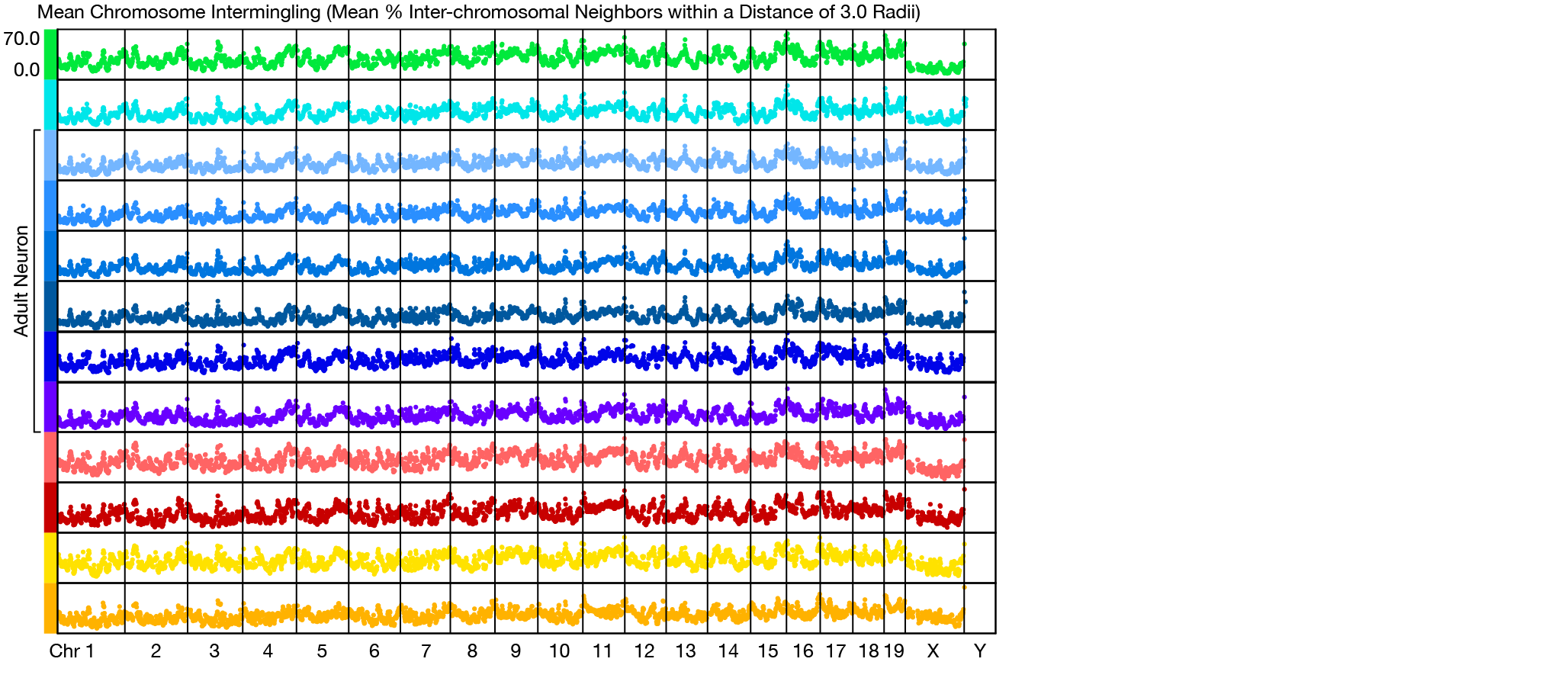

**Figure S10. Mean extent of chromosome intermingling across the genome, for each structure type.**

Similar to Figure 4B, but for the mean extent of chromosome intermingling.

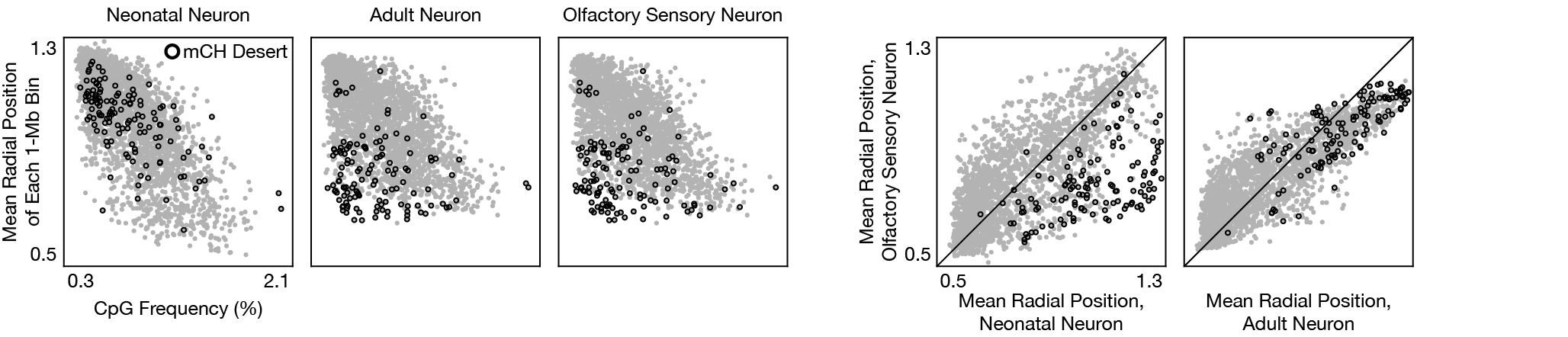

**Figure S11. Comparison of mean radial positions in neonatal neurons, in adult neurons, and in OSNs, and CpG frequency along the genome.**

Similar to Figure 4C left, but plotted against CpG frequency along the genome or mean radial position in OSNs.

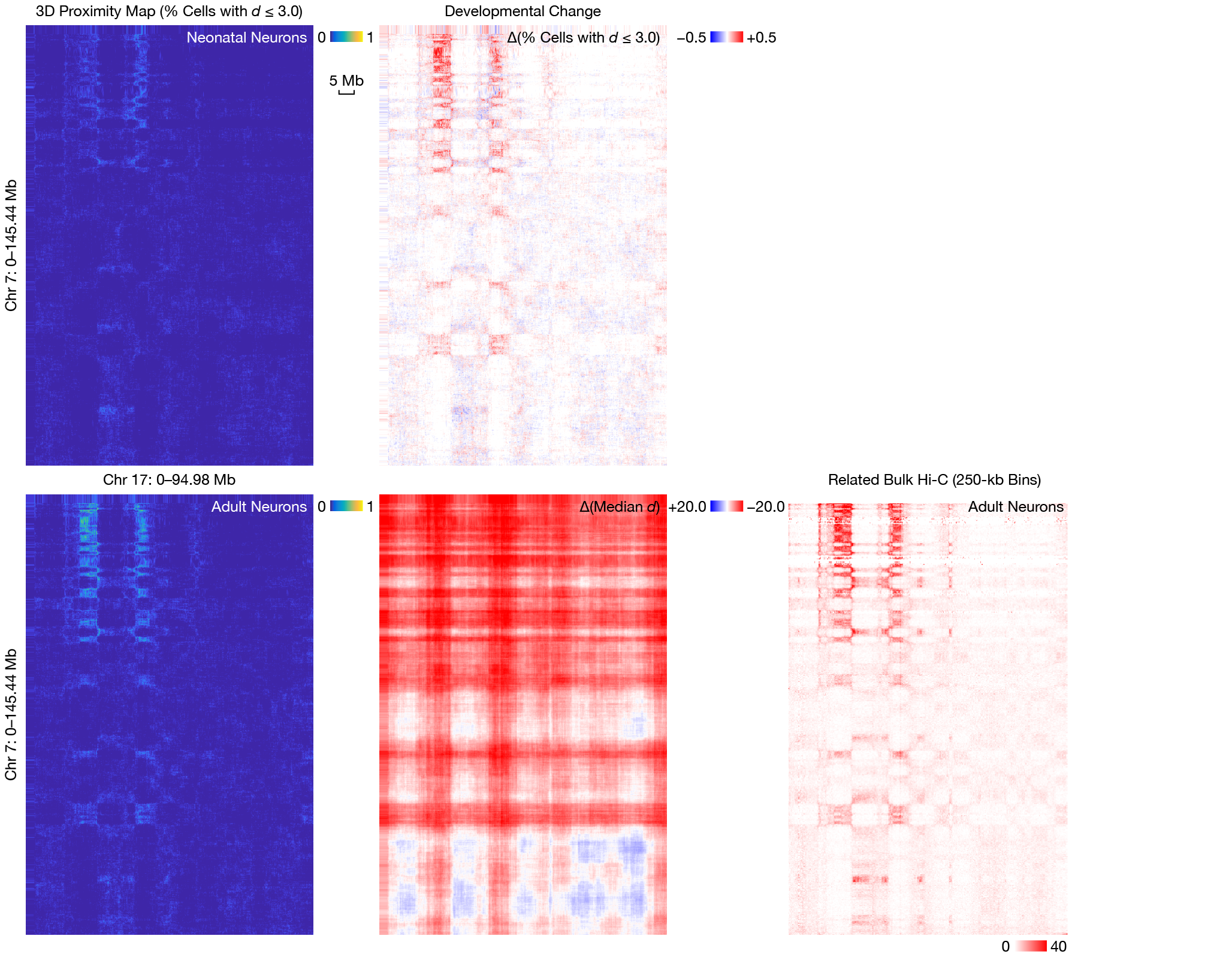

**Figure S12. Developmental change in inter-chromosomal interaction between certain inward-moving regions in neurons.**

Similar to Figure 5C–D, but for inter-chromosomal interaction between the entire Chr 7 and 17. The strongly interacting regions were VR gene clusters.

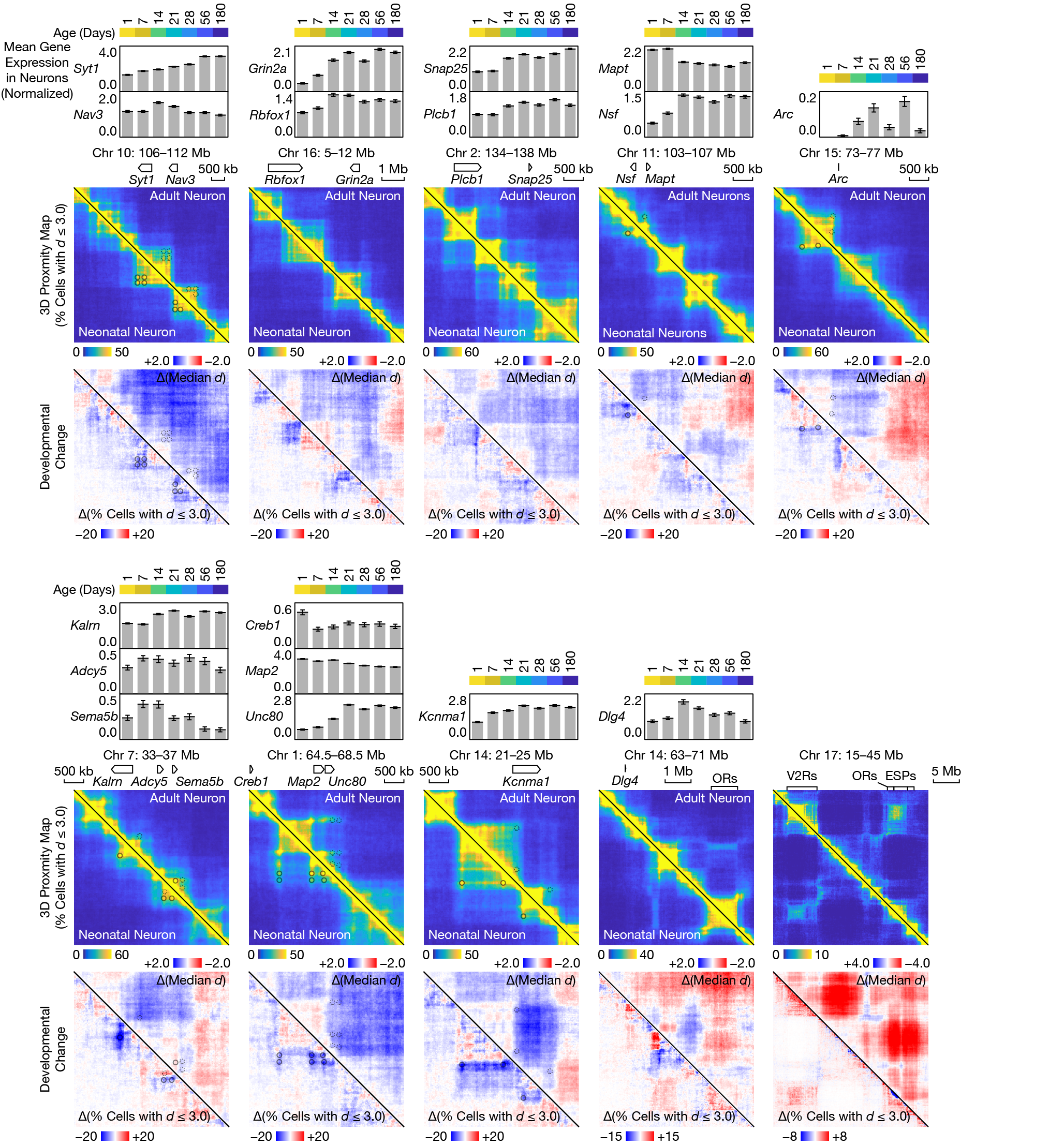

**Figure S13. Developmental change in local 3D genome structure.**

Similar to Figure 5C–D, but for notable changes around other genes critical for synaptogenesis and/or linked to neurodevelopmental disorders.

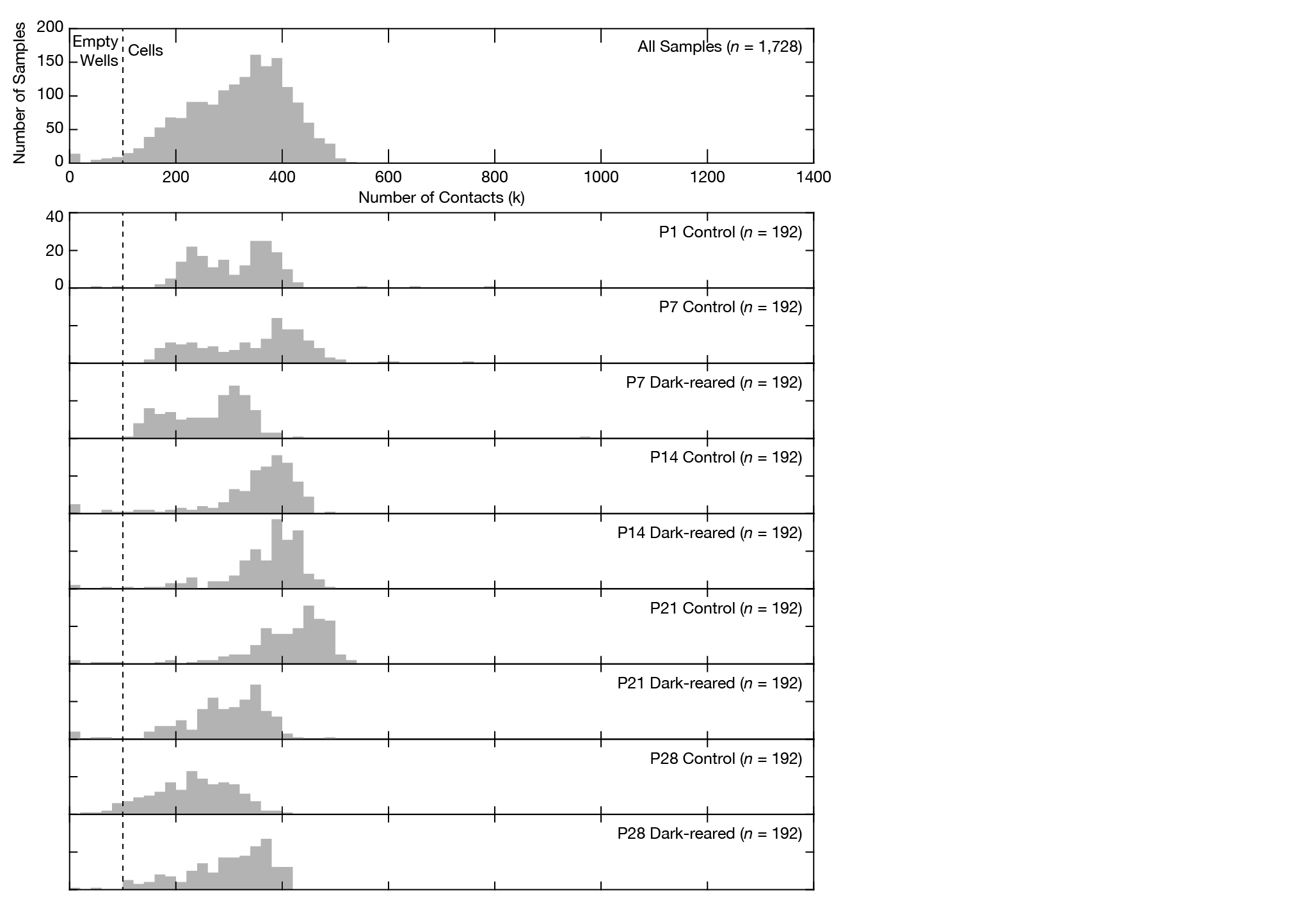

**Figure S14. Histograms of numbers of detected contacts per cell in the sensory deprivation Dip-C dataset.**

Similar to Figure S5, but for the sensory deprivation dataset.

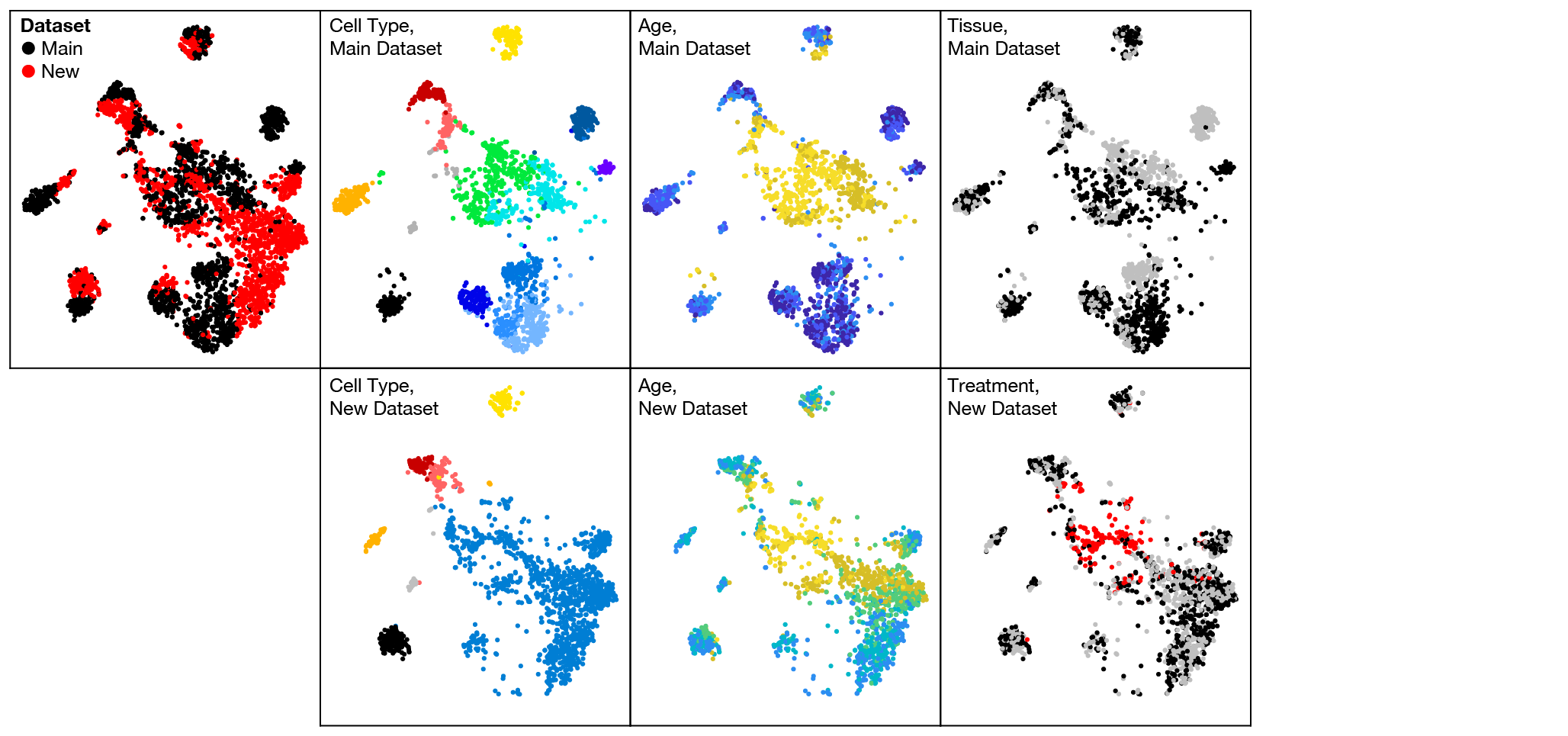

**Figure S15. Joint t-SNE plot of the main and sensory deprivation Dip-C datasets.**

Similar to Figure 3A and Figure 6A, but jointly on the same t-SNE plot. Relative separation of the 2 datasets indicated batch effects from different mouse strains, brain regions, and restriction enzymes.

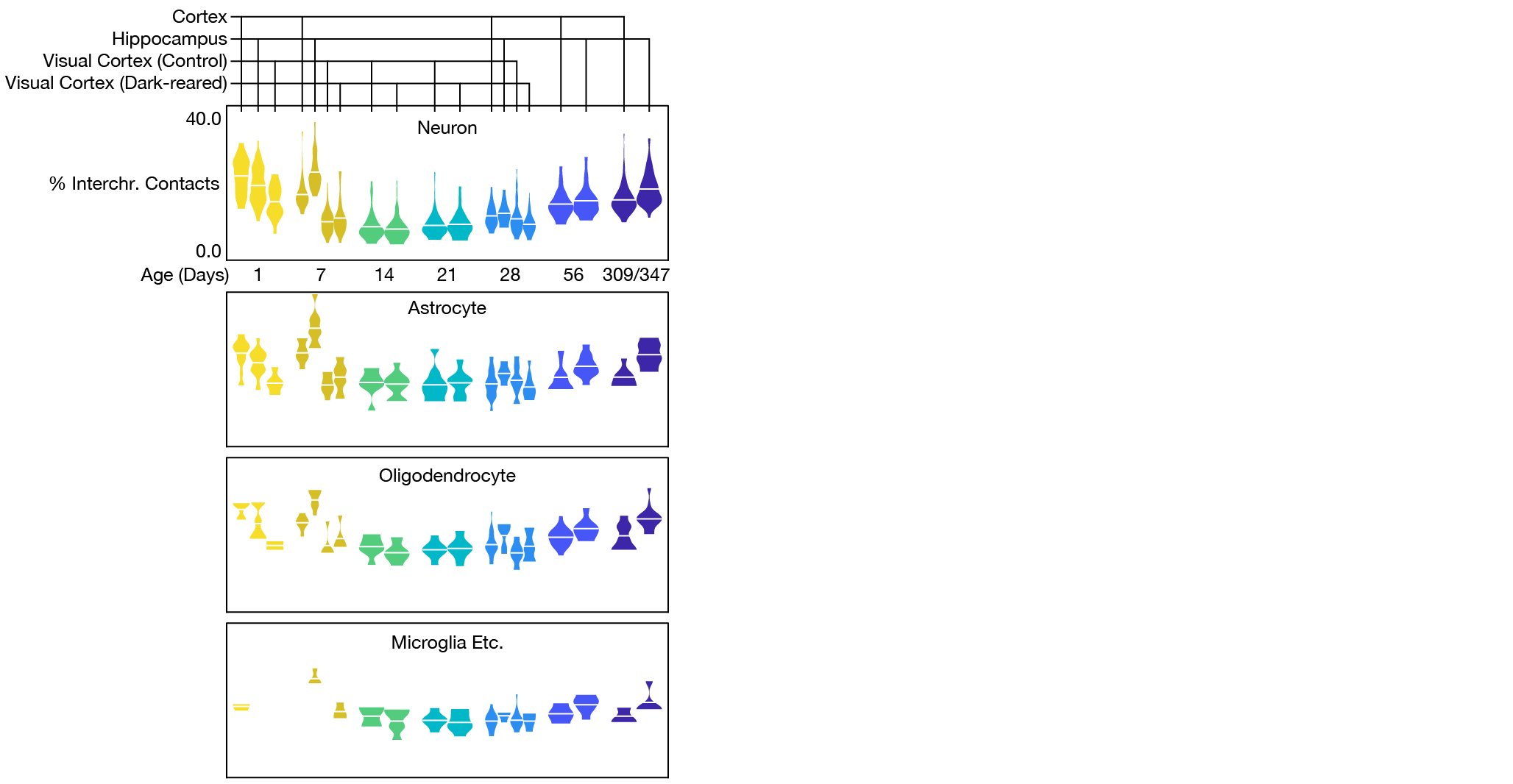

**Figure S16. Percent inter-chromosomal contacts for each sample.**

Similar to Figure 6D, but for percent inter-chromosomal contacts (therefore, the same as Figure 4A bottom).

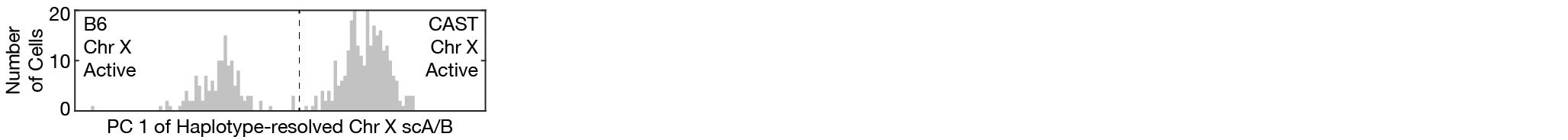

**Figure S17. Histogram of PC 1 of haplotype-resolved Chr X scA/B values.**

The 2 unequal cell populations were consistent with known biased XCI in F1 hybrids.

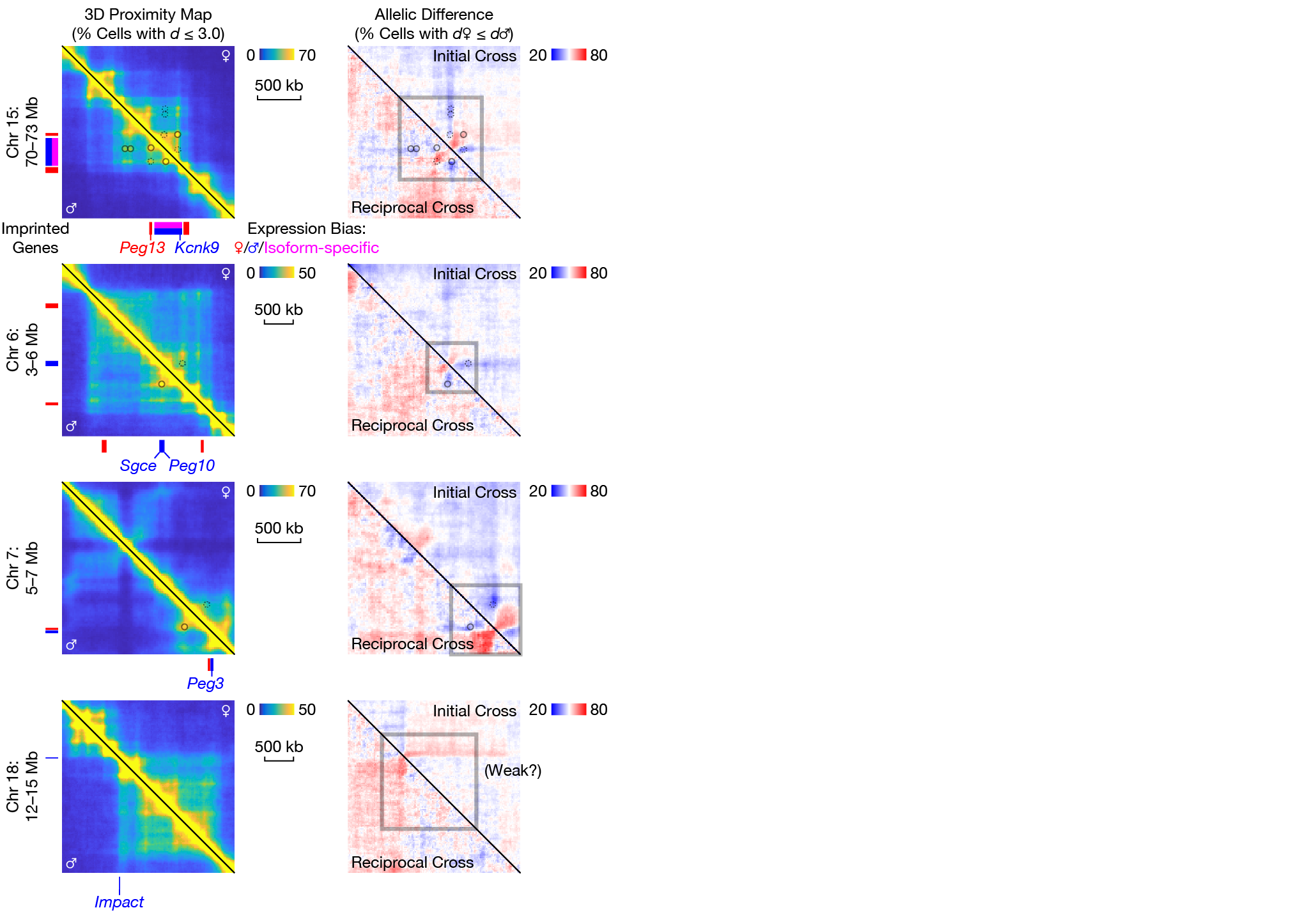

**Figure S18. Parent-of-origin-specific 3D structure around the other 4 imprinted genes/gene clusters.**

Similar to Figure 7C, but for the other 4 genes/gene clusters with parent-of-origin difference.

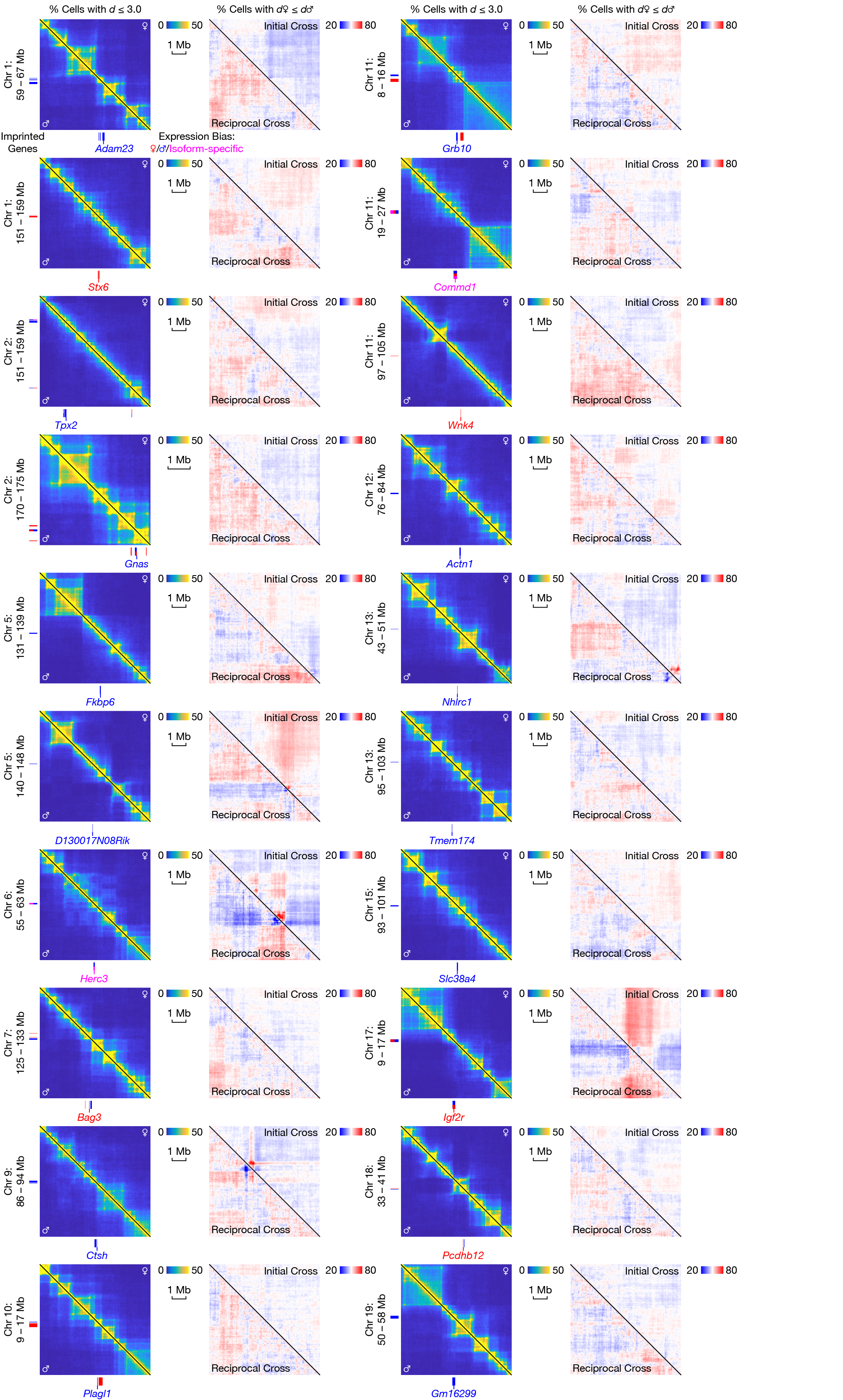

**Figure S19. No detectable parent-of-origin-specific 3D structure around the remaining imprinted genes.**

Similar to Figure 7C, but for the 21 genes/gene clusters without parent-of-origin difference.

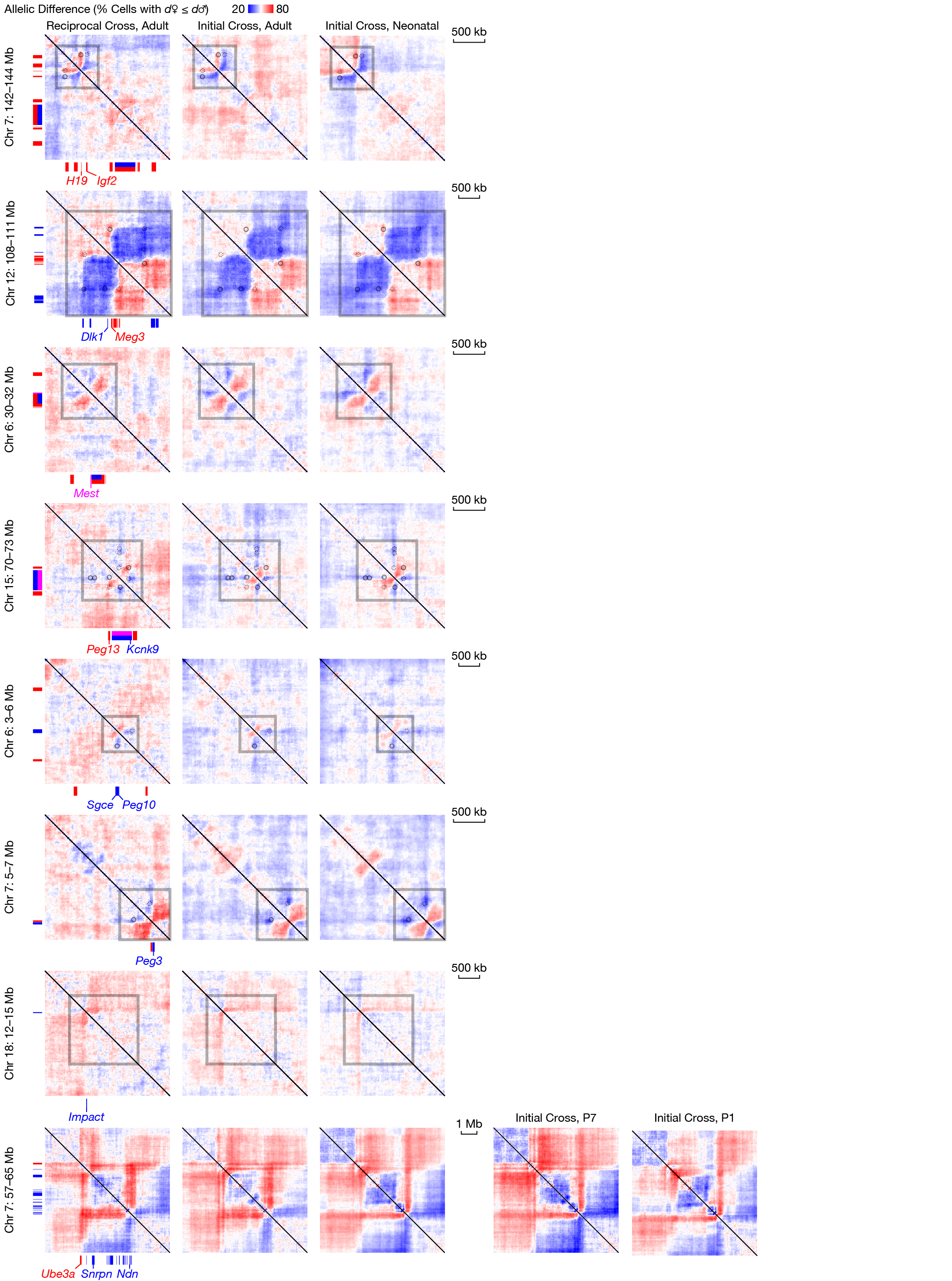

**Figure S20. Parent-of-origin-specific 3D structure around imprinted genes, separately for different ages.**

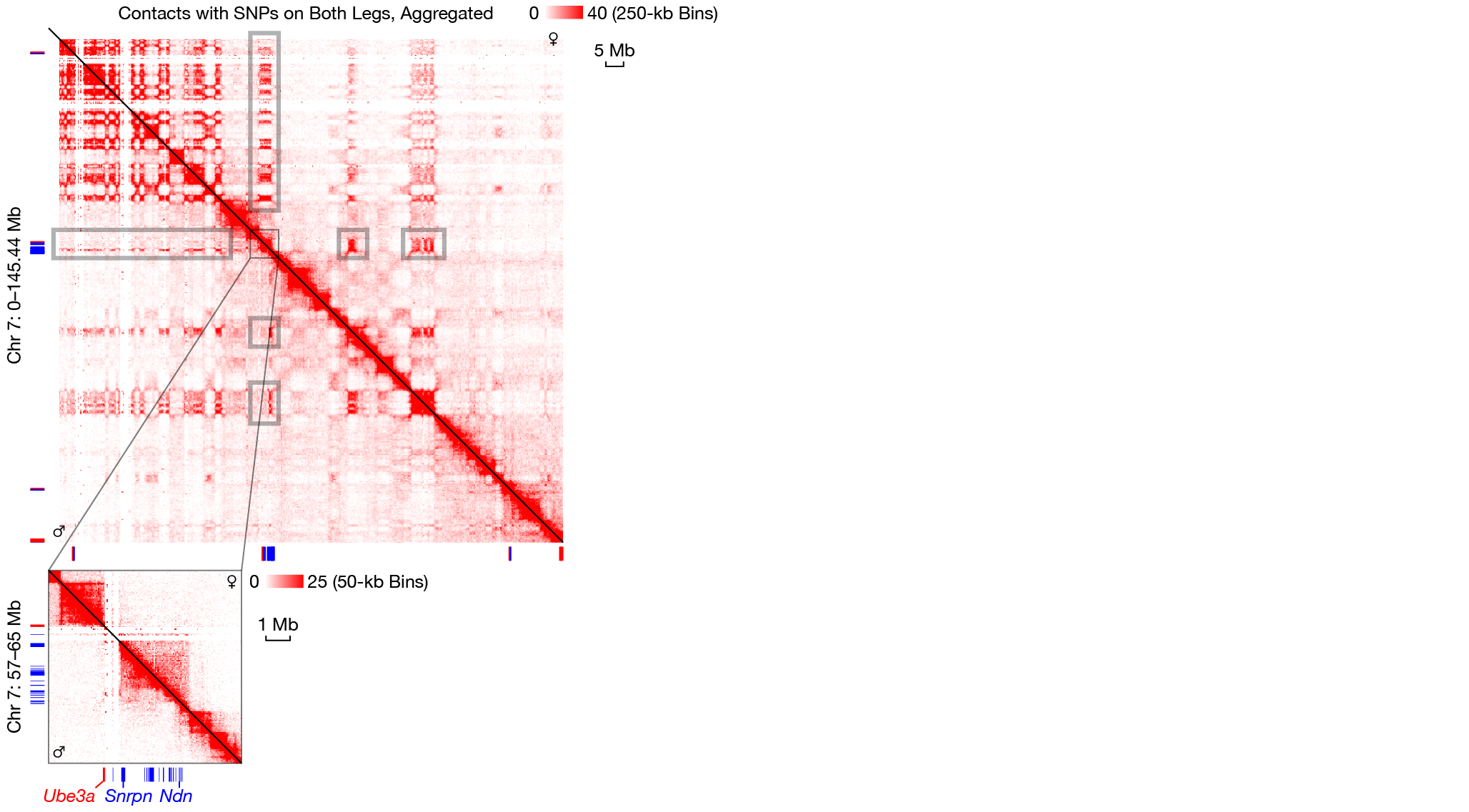

**Figure S21. Parent-of-origin-specific 3D structure at the PWS/AS locus, from the raw contact map.**

No haplotype imputation or 3D modeling was involved.
