## Supplementary Table Titles for "Experience-independent transformation of single-cell 3D genome structure and transcriptome during postnatal development of the mammalian brain"

**Table S1. Cell-type-specific marker genes from MALBAC-DT.**

**Table S2. Cell-type-specific marker genes from MALBAC-DT (among neurons).**

**Table S3. Genes sorted by PC 1 loading from MALBAC-DT (among neurons).**

**Table S4. Genes sorted by PC 1 loading from MALBAC-DT (among neurons; less stringent).**

**Table S5. Neonatal and adult gene modules from MALBAC-DT (among neurons).**

**Table S6. Neonatal and adult gene modules from MALBAC-DT (among neurons; less stringent).**

**Table S7. Information about Dip-C cells.**

**Table S8. Regions with differential scA/B values from Dip-C.**

**Table S9. Cell-type-specific marker genes from published transcriptome data.**

**Table S10. Regions that moved inward during neuron development.**

**Table S11. Genes with developmentally regulated scA/B values or radial positions in neurons.**
