## Supplementary Methods for "Experience-independent transformation of single-cell 3D genome structure and transcriptome during postnatal development of the mammalian brain"

**Animals**

Animal protocols were approved by the Institutional Animal Care and Use Committee (IACUC) at Harvard University and at Peking University. For the MALBAC-DT and sensory deprivation Dip-C datasets, all animals (a pool of 3 pups for P1, and a single animal each for all other samples) were males from the inbred C57BL/6N strain. For the main Dip-C dataset, F1 Hybrids of CAST/EiJ (JAX 000928) and C57BL/6J (JAX 000664) were generated by in-house breeding. The P1 sample (a pool of 2 pups) was female (inferred from sequencing data); all other samples (a single animal each) were male.

**Isolation of cell nuclei for MALBAC-DT**

Cell nuclei were isolated based on protocols from (Krishnaswami et al., 2016; Lacar et al., 2016) with minor modifications. See details below:

Cortex and hippocampus were dissected in ice-cold PBS, and placed in 2 mL nuclei isolation medium with Triton (0.25 M sucrose, 25 mM KCl, 5 mM MgCl2, 10 mM Tris pH 8.0, 1 uM DTT, RNase inhibitor, 0.1% Triton X-100) in a 2 mL Dounce homogenizer (Sigma D8938). Note that DTT concentration was not specified in (Krishnaswami et al., 2016); here we used 1 uM from (Lacar et al., 2016).

Tissues were homogenized with 5 strokes of the loose (A) pestle, and 15 strokes of the tight (B) pestle. The homogenate was centrifuged for 8 min at 100 g, 4 C, and the supernatant removed carefully. The pellet was re-suspended in 1 mL nuclei isolation medium without Triton (0.25 M sucrose, 25 mM KCl, 5 mM MgCl2, 10 mM Tris pH 8.0, 1 uM DTT, RNase inhibitor).

The tube was centrifuged for 8 min at 100 g, 4 C, and the supernatant removed carefully. The pellet was re-suspended in 1 mL nuclei storage buffer (0.1665 M sucrose, 5 mM MgCl2, 10 mM Tris pH 8.0, RNase inhibitor), and filtered with 40 um. Note that sucrose concentration was not specified in (Krishnaswami et al., 2016); here we used 0.1665 M from (Lacar et al., 2016). Also note that the recipe in (Lacar et al., 2016) was inconsistent between ingredient volumes and final concentrations; here we followed the final concentrations.

Nuclei were stained with Hoechst 33342 (except the P14 sample), and sorted with a FACSAria III (BD).

**MALBAC-DT**

MALBAC-DT was performed as previously described (Chapman et al., 2020; Tian et al., 2020).

**Isolation of cell nuclei for Dip-C**

Cell nuclei were isolated based on protocols from (Krishnaswami et al., 2016; Lacar et al., 2016) with minor modifications. In particular, Tris buffer would react with formaldehyde, and was therefore substituted with equal molarities of HEPES buffer. In addition, nuclei were resuspended in the isolation medium, rather than the storage buffer, to better preserve nuclear morphology. See details below:

Cortex and hippocampus were dissected in ice-cold PBS, and placed in 2 mL nuclei isolation medium with Triton (0.25 M sucrose, 25 mM KCl, 5 mM MgCl2, 10 mM HEPES pH 8.0, 1 uM DTT, 0.1% Triton X-100) in a 2 mL Dounce homogenizer (Sigma D8938). Note that DTT concentration was not specified in (Krishnaswami et al., 2016); here we used 1 uM from (Lacar et al., 2016).

Tissues were homogenized with 5 strokes of the loose (A) pestle, and 15 strokes of the tight (B) pestle. The homogenate was centrifuged for 8 min at 100 g, 4 C, and the supernatant removed carefully. The pellet was re-suspended in 1 mL nuclei isolation medium without Triton (0.25 M sucrose, 25 mM KCl, 5 mM MgCl2, 10 mM HEPES pH 8.0, 1 uM DTT).

The tube was centrifuged for 8 min at 100 g, 4 C, and the supernatant removed carefully. The pellet was re-suspended in 1 mL nuclei isolation medium without Triton, and filtered with 40 um.

**Fixation of cell nuclei for Dip-C**

Nuclei were fixed according to our Dip-C protocol (Tan et al., 2019) with modifications. In particular, nuclei were fixed directly in the 1 mL nuclei isolation medium without Triton (with HEPES instead of Tris buffer; see above) rather than PBS, to better preserve nuclear morphology; and centrifugation was performed at 1,000 g rather than 600 g. See detailed below:

Nuclei were fixed directly in the nuclei isolation medium without Triton (see above; 1 mL) with the addition of 66.7 uL 32% PFA (EMS 15714) to a final concentration of 2%. The tube was rotated for 10 min at room temperature. Then 200 uL 1% BSA in PBS was added. The tube was centrifuged for 5 min at 1,000 g, 4 C, and the supernatant removed. The pellet was re-suspended in 1 mL ice-cold 1% BSA in PBS. Nuclei were counted and aliquoted to up to 0.5 million per tube. Each tube was centrifuged for 5 min at 1,000 g, 4 C, and the supernatant removed. The pellet was stored at -80 C.

**Dip-C**

A pellet of fixed cell nuclei was thawed on ice. We then proceeded directly to the step where 50 uL 0.5% SDS was added in the Dip-C protocol (Tan et al., 2019) and followed through.

For the main dataset, nuclei were stained with DAPI, and sorted on a MoFlo Astrios (Beckman Coulter). For the sensory deprivation dataset, nuclei were stained with 7-AAD, and sorted on a FACSAria III (BD). Nuclei were amplified with homemade Nextera. For the main dataset, total of 26.5 96-well plates were processed, leading to 2,544 samples (Figure S5). For the sensory deprivation dataset, a total of 18 plates were processed, leading to 1,728 samples (Figure S14).

For the main dataset, the restriction enzyme was MboI. For the sensory deprivation dataset, the restriction enzyme was 1 U/uL NlaIII (NEB R0125L).

The Dip-C whole-genome amplification procedure was streamlined from 2 aspects. In particular, 12-channel (or 8-channel for Nextera i5 index primers) pipettes were used at all steps; the numbers of index primers were doubled, to 24 i7 primers and 16 i5 primers, allowing simultaneous sequencing of 24 × 16 = 384 samples (4 96-well plates) on the same lane (for example, on a NovaSeq). We have also added a positive control (pure gDNA rather than single cells) for first-time users. A step-by-step protocol, including all reagent catalog numbers and volumes (including overhead), is included as a supplementary protocol.

If assembling one’s own Tn5 transposome is not an option, a roughly equal amount of “TTE Mix V50” (~0.015 uL per cell) from a TD501 kit (Vazyme) can be used in place of homemade Nextera transposome during the transposition step. In this way, the Dip-C procedure (the Nextera version) can be performed entirely with commercially available reagents, without the need for Tn5 transposase production.

**Sensory deprivation by dark rearing**

To deprive the visual system from sensory inputs, mice were reared in complete darkness starting from P1. At desired ages, mice were sacrificed in a dark room under dim red light, until eyes were removed. Visual areas of the cortex were dissected for nuclei isolation.

**Published Data**

*Reference genome.* The mouse reference genome (GRCm38) and gene annotations (ALL) were downloaded from the GENCODE (https://www.gencodegenes.org/mouse/) M19 release.

*Bulk Hi-C data*. For projection onto the t-SNE plot (Figure S8), the first 10 million reads of each data set were downloaded from the SRA with “fastq-dump” (“fastq-dump --split-files --gzip -X 10000000 [SRR accession]”). The accession numbers were: SRR5617733 for adult (3 months) cortical neurons (*NeuN*+) (Jiang et al., 2017), SRR8441362 for adult (3–6 months) hippocampal excitatory neurons (*Camk2a+*) after saline injection (Fernandez-Albert et al., 2019), SRR8441366 for adult (3–6 months) hippocampal excitatory neurons (*Camk2a+*) 1 h after kainic acid injection (Fernandez-Albert et al., 2019), SRR8441370 for adult (3–6 months) hippocampal excitatory neurons (*Camk2a+*) 48 h after kainic acid injection (Fernandez-Albert et al., 2019), SRR5339908 for embryonic (E14.5) cortical neurons (*Hes5-*, *Dcx+*) (Bonev et al., 2017), SRR5339876 for embryonic (E14.5) cortical neural progenitors (*Hes5+*, *Dcx-*) (Bonev et al., 2017), SRR5339832 for *in vitro* differentiated cortical neurons (DIV12+9; *Tau*+, G0/G1 phase) (Bonev et al., 2017), SRR5339786 for *in vitro* differentiated cortical neural progenitors (DIV12+2; *Sox1+*, G0/G1 phase) (Bonev et al., 2017), and SRR941305 for *in vitro* differentiated astrocytes (Sofueva et al., 2013).

For visualization of bulk contact maps, data of *in vitro* differentiated, embryonic-like cortical neurons (“CN”: DIV12+9; *Tau*+, G0/G1 phase) (Bonev et al., 2017) was directly visualized through the Juicebox.js web browser (http://www.aidenlab.org/juicebox/) (Robinson et al., 2018). Note that their *in vivo* embryonic cortical neurons (“ncx CN”: E14.5; *Hes5-*, *Dcx+*) would be a better match, but were not available through the web browser. Data of adult (3 months) cortical neurons (*NeuN*+) was downloaded from the SRA of (Jiang et al., 2017) (accession numbers: SRR5617731 and SRR5617733 for the two replicates, respectively; both male, and originally mapped to the mm9 reference genome), and processed in the same manner as our single-cell data. The resulting “contacts.hic” file was then visualized with Juicebox.js (Robinson et al., 2018).

*Dip-C data.* 3D genome structures of 106 mature OSNs at 20-kb resolution in our previous study (Tan et al., 2019) were downloaded from the GEO (accession number GSE121791, the files “20k.1.clean.3dg”).

*Lists of cell-type-enriched genes and regions.* The following procedure is for Figure 2a. See Table S2 for all gene names.

The top 100 genes in microglia, in myelinating oligodendrocytes, in oligodendrocyte progenitors, and in astrocytes were obtained from the website of (Zhang et al., 2014) (https://web.stanford.edu/group/barres_lab/brain_rnaseq.html; unfortunately no longer available by the time of this submission) by querying the top 100 genes (ranked by fold change) with FPKM ≥ 10 that are enriched in each cell type of interest (versus all other cell types), respectively.

The 10,984 accessible regions in microglia, 18,664 in oligodendrocytes, 17,418 in astrocytes, and 18,470 in SST+ inhibitory neurons were obtained by downloading the differential accessibility file (“atac_matrix.binary.da_results.sig_open.txt”) from the website of (Cusanovich et al., 2018) (http://atlas.gs.washington.edu/mouse-atac/) and separating the peaks based on cell types (“16.3” for microglia, “21.1” for oligodendrocytes, “19.1” for astrocytes, and “15.3” for SST+ inhibitory neurons), respectively.

The top 60 genes up- and down-regulated during astrocyte development (comparing ages P1–8 to P17–30) were obtained from Figs. 6A and 6C, respectively, of (Cahoy et al., 2008).

The 125 genes in hippocampal granule cells, and 108 in (hippocampal) inhibitory neurons were obtained by first downloading Table S1 of (Habib et al., 2016). The TPM values were then back-calculated from the “log TPM values” (which we inferred to mean log_2_ (TPM + 1) based on descriptions in the paper). Finally, enriched genes were identified as TPM ≥ 10 and fold change ≥ 2 between the cell type of interest (“Granule cells DG” and “GABAergic interneurons”, respectively) and each of the 4 other neuronal types.

The 4,896 accessible regions in hippocampal granule cells were obtained by downloading Table S5 of (Lareau et al., 2019) and identifying peaks with their highest values in the cell type of interest (“EN16”).

The 35,230 un-methylated regions in SST+ inhibitory neurons were obtained by downloading Table S5 of (Luo et al., 2017) and navigating to the tab for the cell type of interest (“mSst_1”).

The top 100 genes up-regulated during cortical pyramidal cell development (comparing ages P2 to P11; note that this does not exactly match the time of our 3D genome transformation, but is the closest we can find) were obtained by first downloading the gene expression matrix (“GSM3017261_150000_CNS_nuclei.mat.gz”, in MATLAB format) from the GEO accession (GSE110823) of (Rosenberg et al., 2018). Cortical pyramidal cells were then extracted by their cell-type clusters (“cluster_assignment” containing “CTX Pyr” or “CLAU Pyr”; see Figure 3C of (Rosenberg et al., 2018)), and separated by age (“sample_type” being “p2_brain” or “p11_brain”). Expression values (“DGE”) were normalized into TPM values (so that each cell sums to 1,000,000), and averaged among P2 and P11 cortical pyramidal cells, respectively. Finally, up-regulated genes were identified as the top 100 genes (ranked by fold change between P11 and P2) with TPM ≥ 100 in P11 cells. Note that similar to our previous observation in olfactory sensory neurons (Tan et al., 2019), the top developmentally down-regulated genes on average did not show changes in single-cell A/B compartment values.

Outdated gene names were converted to modern ones by looking up the Mouse Genome Informatics (MGI) website (http://www.informatics.jax.org/).

*mCH desserts.* The following procedure is for Figure 3c and Extended Data Figs. Genomic coordinates of mCH desserts were obtained by downloading Table S2 of (Lister et al., 2013), navigating to the mouse section, and lifting over from mm9 coordinates to mm10 ones with the online LiftOver tool of the UCSC Genome Browser (https://genome.ucsc.edu/cgi-bin/hgLiftOver; parameters were: “Minimum ratio of bases that must remap” = 0.01 and “Allow multiple output regions”, because of the repetitive nature of these regions).

*Imprinted genes.* The following procedure is for Figure 4 and Extended Data Figs. Imprinted genes in the mouse brain (cerebellum), their genomic coordinates, and their expression biases were obtained from tab “(G)” of Supplementary File 1 of (Perez et al., 2015).

**Data Analysis for MALBAC-DT**

*Generation of the count matrix.* Raw data were pre-processed as previously described (Chapman et al., 2020; Tian et al., 2020) to generate a matrix of UMI counts for different genes and cells.

*Normalization.* In RStudio, the count matrix was loaded into Seurat v3 (Stuart et al., 2019) with “CreateSeuratObject”, and normalized with “SCTransform” (Hafemeister and Satija, 2019), which by default kept the top 3,000 variable genes for some subsequent analysis (including PCA, UMAP, clustering, but not including marker identification).

*PCA.* In Seurat, PCA was preformed with “RunPCA”. For neurons, percent variance explained by each PC was calculated by dividing the variance of each PC (obtained from the Seurat object with “@reductions$pca@stdev”, followed by squaring) by the total variance (of the 3,000 variable genes; obtained from the Seurat object with “@reductions$pca@misc$total.variance”). PC 1 score of each cell was obtained with “Embeddings”. PC 1 loading of each variable gene was obtained with “Loadings”.

*UMAP.* In Seurat, transcriptome types were visualized with “RunUMAP” (with parameters “dims = 1:15” for all cells, and “dims = 1:30” for neurons).

*Identification of transcriptome types by Louvain clustering.* In Seurat, raw cell-type clusters were identified with “FindNeighbors” (with parameters “dims = 1:15” for all cells, and “dims = 1:30” for neurons, the same as in UMAP) and “FindClusters” (with default paramters for all cells, and “resolution = 4” for neurons). Final transcriptome types were then generated by manually merging raw clusters based on expression patterns of known marker genes, with “RenameIdents”.

*Isolation of neurons for detailed analysis.* As recommended by Seurat tutorials, neurons were separately analyzed in detail. In particular, a new Seurat object was created with “subset”; all analysis—starting from “SCTransform”—were performed again on the new object.

*Identification of cell-type-specific marker genes.* In Seurat, raw marker genes specific to different cell populations were identified with “FindMarkers” (with parameters “only.pos = TRUE, min.pct = 0.25, logfc.threshold = 0.25”, as suggested in tutorials on the Seurat website, “ident.1” specifying the cell population of interest, and “ident.2” specifying the background population). Among raw marker genes, the top 100 were then selected with “%>% top_n(n = 100, wt = avg_logFC)”, as suggested in Seurat tutorials.

*Identification of correlated gene modules.* Normalized expression matrix of the top 3,000 variable genes was exported from Seurat with “t(as.matrix(GetAssayData(seurat_object_neuron)))[,VariableFeatures(seurat_object_neuron)]”. Correlated gene modules were identified with WGCNA (Langfelder and Horvath, 2008) according to tutorials on its website. In particular, the soft-thresholding power was first chosen to be 6, after visualizing the output of “pickSoftThreshold” (with parameters “corFnc = "bicor", networkType = "signed"”). Modules were then identified with “blockwiseModules” (with parameters “power = 6, corType = "bicor", networkType = "signed", pamRespectsDendro = FALSE”). The 2 largest modules, named “turquoise” and “blue” by WGCNA, were denoted the neonatal and adult modules, respectively.

Expression of module eigengenes in each cell was extracted with “unlist(wgcna_modules$MEs[paste0("ME", "turquoise")])”.

We also generated a less stringent set of gene modules (Table S6), by keeping the top 10,000 variable genes (rather than 3,000 by default) during normalization. In particular, “SCTransform” was performed with parameters “variable.features.n = 10000”. In this case, patterns of percent variance explained by each PC (Figure 2C) remained similar, although now PC 1 explained 2.3% of the total variance (rather than 3.6%). In WGCNA, the soft-thresholding power was chosen to be 8, after visualizing the output of “pickSoftThreshold” (with parameters “corFnc = "bicor", networkType = "signed", blockSize = 10000”). Modules were identified with “blockwiseModules” (with parameters “power = 8, corType = "bicor", networkType = "signed", pamRespectsDendro = FALSE, maxBlockSize = 10000”). The 2 largest modules, named “blue” and “turquoise”, were denoted the neonatal and adult modules, respectively. Note that because of the minor procedural difference, gene modules under these less stringent criteria did not include every single gene from the normal criteria (for example, *Zbtb20*).

**Data Analysis for Dip-C**

*Generation of contact maps.* Single-cell contact maps were generated as we previously described (Tan et al., 2019), with “hickit” and “dip-c” packages.

*Filtering out empty wells.* In the main dataset, out of all 2,544 samples (26.5 96-well plates), we removed 590 (23%) that yielded < 100 k contacts per sample (Table S7, Figure S5). Similarly, in the sensory deprivation dataset, 36 (2%) out of 1,728 samples (18 plates) were removed (Table S7, Figure S14). Most of these “empty samples” were wells missed by the flow cytometer.

*Single-cell A/B compartment (scA/B) values.* scA/B values were calculated as we previously described, from contact maps with “dip-c color2” (with parameters “-b1000000 -H -c color/mm10.cpg.1m.txt” for 1-Mb bins, or “-b100000 -H -c color/mm10.cpg.100k.txt” for 100-kb bins in Figure S2) (Tan et al., 2019).

Only autosomal bins that were present in all cells were retained. scA/B values were rank-normalized to 0–1 in each cell with MATLAB “tiedrank” (“(tiedrank(compartment_values_in_cell)-1)./(num_bins_in_cell-1)”).

After PCA with MATLAB “pca” (“pca(compartment_value_matrix)”), only the first 20 PCs (“pca_score(:, 1:20)”) were retained for further visualization and clustering.

t-SNE was performed with MATLAB “tsne”, initialized by the first 2 PCs (“tsne(pca_score(:, 1:20), 'InitialY', pca_score(:, 1:2))”). UMAP (Figure S2) was performed with MATLAB “run_umap” (“run_umap(pca_score(:, 1:20))”).

Hierarchical clustering was performed with MATLAB “linkage”, on Euclidean distances of the first 20 PCs with Ward’s method (“linkage(pca_score(:, 1:20), 'ward', 'euclidean')”). Similar to transcriptome analysis (see above), the 13 major and 3 minor (unknown) structure type clusters were then manually merged from raw clusters with the t-SNE plot as a visual aid.

*Projection of bulk Hi-C data onto the t-SNE space.* The first 10 million reads of each bulk Hi-C data were processed in the same manner as our single-cell data, and projected onto the PCA space as we previously described (Tan et al., 2019), with MATLAB. Subsequently, because additional data points cannot be directly projected onto an existing t-SNE space, the final plot (Figure 1c right) was generated by performing t-SNE jointly on both bulk and single-cell data with MATLAB “tsne”.

*Integrative analysis of scA/B values, transcriptomes, methylomes, and chromatin accessibility.* For each gene or region of interest, the 1-Mb bin to which its mid-point belonged (calculated according to GENCODE for genes) was first identified. The same filtering as above (only autosomal bins that were present in all cells were retained) was then applied.

For each list of genes (Table S1 and Table S2 from our MALBAC-DT dataset, or Table S8 from published data) or regions, normalized scA/B values of the above bins were averaged in each cell, producing a single-cell, list-averaged value. Note that for some lists, each bin might be counted more than once (especially for accessible or un-methylated regions, because there were only ~3,000 1-Mb bins across the genome); in this case, the average scA/B value was a weighted average of bins.

The distribution of these average scA/B values were visualized either as a heatmap (Figure 3D, Figure S9) or on the t-SNE plot (Figure S7).

*Identification of regions with differential scA/B values.* To systematically identify regions with differential scA/B values between specific cell populations (Table S8, Figure 3E), we compared scA/B values of each 1-Mb bin between two cell populations with two-sided Mann–Whitney U tests (“ranksum” in MATLAB), in a manner similar to marker gene identification in transcriptome studies (see above). FDR was calculated with the Benjamini–Hochberg procedure (“mafdr” in MATLAB with parameters “'BHFDR', true”).

For each cell population, the top 100 regions that changed scA/B by at least 0.03 (separately for increase and decrease), sorted by P-values (thus FDR), were identified. Genes whose mid-points fell into each region were further identified.

*Basic characteristics of single-cell contact maps.* For each cell, the extent of chromosome intermingling was quantified as the fraction of inter-chromosomal contacts (Tan et al., 2019) (Figure 4A bottom). Here we calculated this fraction from raw contacts before removal of duplicates, which has the advantage of not depending on sequencing depths, and differs only slightly from that in duplicate-removed contacts. To avoid sex bias, we analyzed only autosomal contacts.

*Generation of 3D structures.* Single-cell 3D genome structures were generated at 20-kb resolution as we previously described (Tan et al., 2019), with “hickit” and “dip-c” packages.

We generated 5 replicate structures per cell with different random seeds (1–5). After the same filtering as we previously described (the bottom 6% of 20-kb particles that harbored the least numbers of haplotype-resolved contact legs within ±0.5 Mb were removed; these regions were typically unmappable and thus difficult to model) (Tan et al., 2018; Tan et al., 2019), a root-mean-square (across pairs of replicates) root-mean-square (across 20-kb particles) deviation (r.m.s.d.) value was calculated for each cell. Only cells with r.m.s.d. ≤ 1.5 particle radii (high-quality 3D structures) were retained for further analysis.

*Radial positioning.* The radial preference of each 1-Mb bin was calculated as we previously described (Tan et al., 2019), with “dip-c color -C”. Note that this value (3D distance to the nuclear center of mass) was not on a uniform scale, because a spherical shell of the same thickness contained more chromatin (larger volume) at a larger radius.

*Changes in 3D genome structure during postnatal neuronal development.* We visualized pairwise 3D chromatin interactions between 20-kb particles by first calculating a matrix of pairwise 3D distances within a genomic region (or between two regions) in each cell with “dip-c pd” (“dip-c pd -1 region.leg” within one region; or “dip-c pd -1 region_1.leg -2 region_2.leg” between two regions, such as inter-chromosomal analysis in Figure S4a). The two alleles were calculated separately.

A 3D proximity map—a matrix of fractions of cells where each pair of 20-kb particles are nearby in 3D (distance ≤ 3.0 radii) (for example, Figure 3d left) (Tan et al., 2019)—was then generated by binarizing each single-cell matrix of pairwise 3D distances and averaging them, with “scripts/threshold_np_float.py” of the “dip-c” package (“scripts/threshold_np_float.py 3”).

Separately, a matrix of median 3D distances between each pair of 20-kb particles was generated by calculating element-wise medians, with “scripts/median_np_float.py” of the “dip-c” package.

Developmental changes of the two above matrices were calculated by subtracting the neonatal matrix from the adult one, with “scripts/subtract_np_float.py” of the “dip-c” package.

*Chromosome intermingling in 3D structures.* The extent of chromosome intermingling of each 1-Mb bin (Figure S5) was calculated as we previously described (Tan et al., 2019), with “dip-c color -i3” (note than this calculates the fraction of intra-chromosomal, or intra-homologous to be exact, 3D neighbors; the fraction of inter-chromosomal neighbors is 1 minus this value).

*Parent-of-origin-specific 3D genome structure.* Similar to our developmental analysis (see above), matrices of fractions of cells where each pair of 20-kb particles are nearby in 3D (distance ≤ 3.0 radii) and of median 3D distances between each pair of 20-kb particles were generated with “dip-c pd”, “scripts/threshold_np_float.py”, and “scripts/median_np_float.py”, but for the maternal and paternal alleles (for example, Figure 3c left), and for parent-of-origin analysis, further separated between the initial and reciprocal crosses.

Within the initial and reciprocal crosses, respectively, allelic differences (for example, Figure 3c middle) were calculated with “scripts/subtract_np_float.py” of the “dip-c” package.

For representative 3D structures (Figure 3c right), the region of interest was extracted from the full 3D genome file (“20k.1.clean.3dg”) with “dip-c reg3” (“dip-c reg3 -i region.reg”), and converted to an mmCIF file with “dip-c color” and “dip-c vis” (“dip-c color -l color/mm10.chr.len” and “dip-c vis -c”) for visualization in PyMol.

Haplotype-resolved raw contact maps (Figure S9) were generated by extracting contacts with haplotype-informative SNPs on both legs (both “0” phases for paternal, both “1” for maternal) from contact maps (“contacts.pairs.gz”) before any haplotype imputation (before “hickit -u”).

**GO enrichment analysis**

GO enrichment analysis of gene lists were performed with the Gene Ontology website (http://geneontology.org/, setting organism to “Mus musculus”) with default parameters. This function was provided by PANTHER (Thomas et al., 2003). Note that the output included both enriched (“+”) and depleted (“−”) terms. Terms were sorted by FDR in Supplementary Tables.

**Data Availability**

Raw sequencing data were deposited at the National Center for Biotechnology Information with accession number PRJNA607329 at https://www.ncbi.nlm.nih.gov/sra/PRJNA607329. Processed data, including files for interactive viewing in the Juicebox web browser, were deposited at with GEO Series accession number GSE146397 at https://www.ncbi.nlm.nih.gov/geo/query/acc.cgi?acc=GSE146397.

**Code Availability**

Code is available at GitHub (https://github.com/tanlongzhi/dip-c and https://github.com/lh3/hickit).
